# How can noise alter neurophysiology in order to improve human behaviour? A combined tRNS and EEG study

**DOI:** 10.1101/2020.01.09.900118

**Authors:** James G. Sheffield, Gal Raz, Francesco Sella, Roi Cohen Kadosh

## Abstract

Random noise has been shown to improve the detection of suboptimal signals in humans and machines. Based on that, transcranial random noise stimulation (tRNS) has aimed to improve human behaviour by targeting neuronal activity. To uncover the poorly understood mechanistic underpinnings of tRNS, we recorded electroencephalography data during arithmetic training while delivering active or sham tRNS above the dorsolateral prefrontal cortex (dlPFC). By successfully removing the tRNS artefact in the time and frequency domains, we examined the mechanisms that underlie its behavioural improvement. We found that active tRNS improved arithmetic performance and impacts specific ERPs components that are associated with attentional mechanisms. Furthermore, the tRNS effect was maximal in individuals with suboptimal arithmetic ability and neurophysiological measures of top-down control and excitation/inhibition ratio. These results providing a novel mechanistic explanation for the effect of tRNS on human behaviour and highlight how suboptimal task-specific behaviour and neurophysiology predicts its effect.

## 1. Introduction

The first association that most people have with the word “noise” is negative. Indeed, in an ideal system, random noise is disruptive. Yet, in non-ideal and non-linear systems, such as the brain, noise can be beneficial, and this has been demonstrated across a number of research fields, including perception, ecology, and engineering^1^. In all of these cases, when noise is applied to a subthreshold signal/input it will improve performance/output. For example, in visual perception, when a stimulus is presented with poor visibility, the addition of an optimal amount of noise can improve an individual’s detection performance^2^. Similar effects have been also observed in the auditory system^3,4^.

This phenomenon has led to the hypothesis that directly adding noise at the neural level, rather than to the stimulus, may confer cognitive benefits. In line with this view, an increasing body of literature highlights the successful application of transcranial random noise stimulation (tRNS), a form of transcranial electrical stimulation, to modulate performance in a variety of training paradigms and cognitive tasks^5,6^. tRNS has been used to improve perception ^7–9^, numerosity discrimination^10,11^, executive control and multitasking^12^, motor skill learning^13,14^, fluid intelligence^15^, and arithmetic learning^16–18^. Furthermore, tRNS has been reported in several studies to yield a stronger effect than anodal transcranial direct current stimulation^6,7,19,20^. Combined, these studies suggest that tRNS is a promising form of neuromodulation for improving complex cognition. Moreover, its higher perceptual threshold compared to other forms of tES makes it easier to create viable sham conditions, as higher amplitudes are required to detect the application of stimulation^21^.

Despite the consistent application of tRNS to alter different cognitive functions, our understanding of its underlying neurophysiological mechanism is poor. Magnetic resonance-based methods are less optimal, especially in the context of cognitive training, due to the induced noise, limited ecological validity, time-constraints, and cost. In contrast, as concurrent stimulation-electroencephalography (EEG) systems become more ubiquitous, the ease-of-use, cost efficiency, and extensive literature makes EEG a promising method for understanding tRNS-induced neural changes. Largely, studies that use EEG to investigate the changes evoked by tRNS have done so by looking at pre/post changes in brain activity. To a degree, this reflects difficulties in simultaneously recording the electrical activity from the brain whilst delivering electrical stimulation, which is often done at amplitudes an order of magnitude larger than the physiological signal measured by EEG equipment. However, one cannot conclude that the changes that follow from the end of the stimulation (termed “offline effect”) are reflective of the altered neural response during stimulation itself (termed “online effect”). An effective procedure to overcome these difficulties could help elucidate possible mechanisms responsible for tRNS-evoked changes.

In a first step to solve this issue, Rufener et al.^22^ used frequency band-pass filters to remove tRNS-induced artefacts from EEG data analysed in the time domain, revealing reduced N1 latency in the tRNS group compared to the sham during a pitch discrimination task. Whilst this method allows demonstration of changes in the time domain, EEG data are multidimensional; we can characterise it not just in the time domain, but also in various facets of the frequency domain, including power, phase, and frequency. Additionally, as the process of creating an event-related potential (ERP) acts as a low-pass filter, it mitigates tRNS noise in higher frequency bands, and though tRNS is usually applied at higher frequency ranges (e.g., 100–640Hz), this process does not remove low-frequency artefacts.

Removing stimulation noise from ongoing EEG activity will allow us to investigate neurophysiological changes that are associated with the effect of tRNS on behaviour, including the causal mechanisms in altering cognition and learning. This step is paramount as it will increase the confidence in the behavioural results and allow improving the tRNS efficacy even further. A prevalent hypothesis is that the effect of tRNS on behaviour is due to stochastic resonance^2,19^, a phenomenon in which introducing an appropriate level of noise can enhance the output of subthreshold signals. Some studies lend support to the stochastic resonance explanation^2–4,18^, and some evidence for stochastic resonance is based on the beneficial effect of tRNS amongst participants with poorer, rather than stronger, baseline ability^12,23,24^. However, the support for the stochastic resonance explanation of tRNS is limited to the behavioural level, and, at least in some cases, might arise from a ceiling effect due to high performance in easy tasks or high-performing participants.

While the neurophysiological signals that tRNS is working on are poorly defined, it was shown that neurophysiological measures before applying tRNS could predict whether tRNS leads to improvement. For example, a previous study found that 1mA tRNS efficacy on sustained attention was dictated by participants pre-stimulation resting theta/beta ratio (TBR)^25^, a measure of top-down control the dlPFC has over midbrain structures such as the anterior cingulate cortex, amygdala, and thalamus^25^. Specifically, participants who were high in TBR, and thus more prone to attentional failing^25,26^, were more likely to show improved sustained attention following 1mA tRNS. By comparison, tRNS in participants with low TBR, who typically demonstrate better sustained attention without tRNS intervention, demonstrated no improvement. Potentially, the improvement that tRNS confers in varied contexts of cognitive tasks, as we described earlier, might be rooted in sustained attention. For example, when tRNS is applied during an arithmetic training, participants’ learning heavily depends on sustained attention^27,28^. Therefore, TBR may predict the effect of tRNS by improving sustained attention during tRNS. Alternatively, TBR might reflect the underlying background noise (the aperiodic signal) of the power spectrum^29^. High aperiodic exponents are thought to reflect lower excitation/inhibition ratios (E/I ratio)^30,31^. Given that tRNS is thought to increase excitation in stimulated sites^32–34^, it is expected that primarily participants with low E/I ratios would benefit from transcranial stimulation^35,36^.

Here, we present the results of a single-session mathematical training study in which we recorded EEG during active or sham tRNS while participants solved complex multiplication problems. The motivation to use mathematical training was threefold: 1) math represents a realistic mode of complex cognition due to its requirement on cognitive processes such as sustained attention and executive functions^27^, emotional regulation^37^, spatial abilities^38^, and algorithmic and memory processes^39^; 2) math is a strong predictor of academic performance in childhood and adolescence^40^, individual outcome, and job performance^41^. Therefore, improving this cognitive function can have important societal implications; 3) most of the existing tRNS studies at the cognitive domain have focused on improving mathematical performance. However, the mechanisms for such enhancement have remained unclear^6^. We tested whether we could successfully remove the artefact induced by tRNS from the ongoing EEG data in both the time and frequency domains. We then used the tRNS cleaned data to investigate the neurophysiological mechanisms that are involved in tRNS enhancement and their effect on behaviour.

## 2. Methods

### 2.1. Pre-registration

This study was pre-registered on Open Science Framework prior to being run (https://bit.ly/2YBITZF). The analyses investigating the effects of stimulation on behavioural outcomes during the task (reaction time [RT], accuracy, and calculation/retrieval analysis) as well as the effects of resting state TBR on task behavioural measures were preregistered. The online analysis of EEG data was not preregistered as we had not yet tested whether tRNS noise could be removed from the EEG data.

### 2.2. Participants

Eighty-one participants (44 females, mean age=23.7 years, SD=5.03) were asked to complete a two-day arithmetic learning paradigm in a study investigating the effects of tRNS stimulation on learning. Participants were excluded if they had a previous history of neurological disorders, a family history of seizure disorders, metal implanted in their head, had previously undergone a neurosurgical procedure, or had received any other form of brain stimulation in the past week. Participants were also requested to refrain from alcohol (for the past 24 hours) or caffeine (in the hour leading up to the study), and to have gotten at least 6 hours of sleep the night before their testing sessions. Four participants were removed during the collection process: one due to software failure, and three due to not returning for a second day of testing. Two participants were removed from behavioural analysis due to having RT scores greater than 3 standard deviations from the mean. During EEG preprocessing analysis, 7 participants were removed because they did not provide enough clean EEG data for analysis, and one participant had a corrupted EEG file. Two of the participants removed this way overlapped with those removed from the behavioural analysis. Final comparisons for the behavioural data were carried out on 75 participants, and the EEG analyses were carried out on 69 participants.

### 2.3. Arithmetic learning task

In each trial, participants were presented with a fixation cross (1000ms) before seeing a multiplication problem, which was left up until the participants responded. Participants responded via a microphone in front of them to measure RTs, and then via a keyboard to evaluate correctness. If participants realised their answer was wrong following their verbal response but before their typed response, they were asked to type in their verbal response anyway. Following their typed response, they received feedback for 2000ms (**Figure 1**).

**Figure 1:**
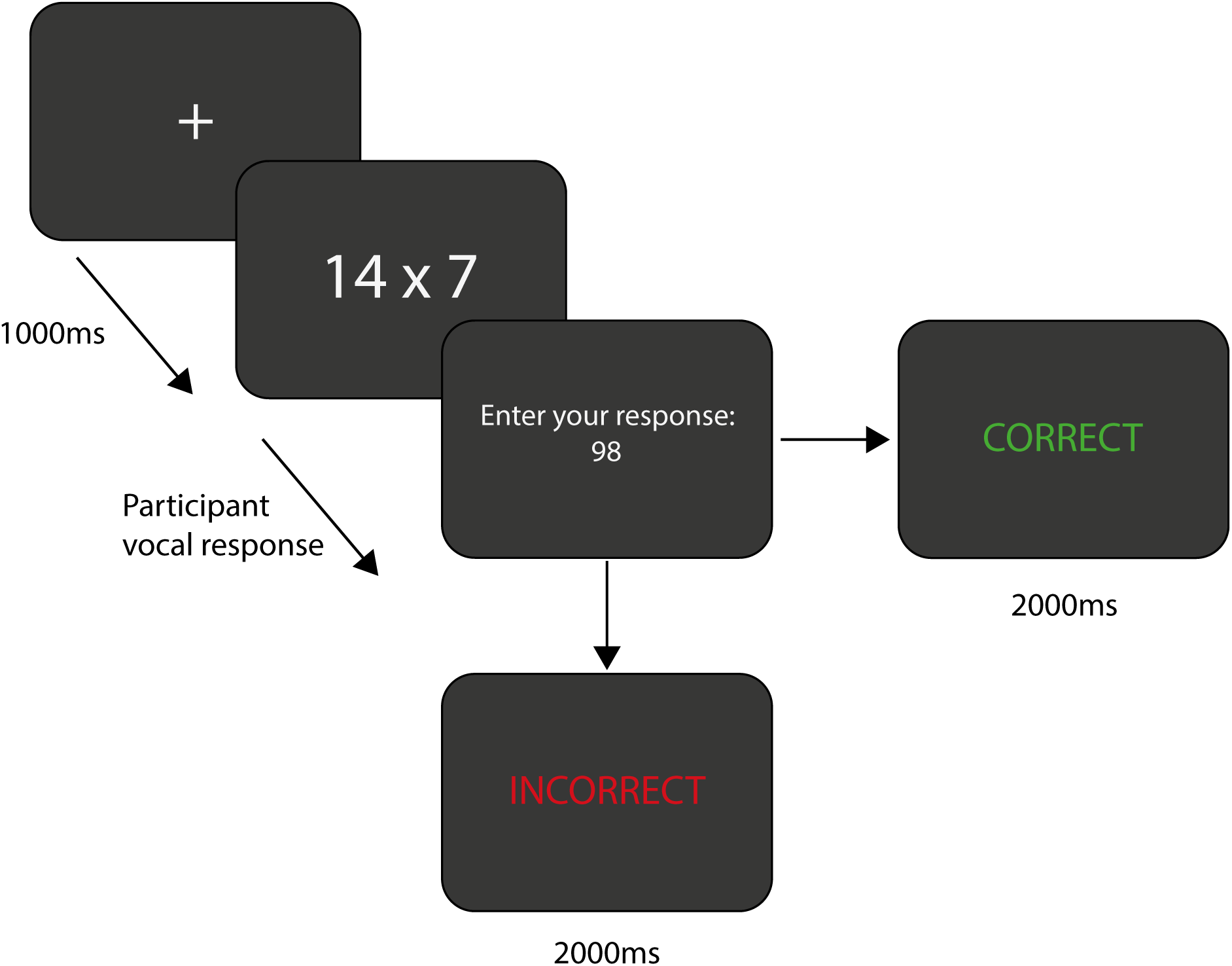
An illustration of a trial from the arithmetic training task. A fixation cross appeared for 1000ms, followed by a multiplication problem that was presented until the participant responded vocally. They were then asked to type their response. Following their response, a feedback was provided for 2000ms

Each multiplication was a two-digit number multiplied by a one-digit number resulting in a two-digit number (see **Table S1**). Neither of the operands included a 0 or a 5, nor were there operands with identical digits. Participants completed 18 blocks, and each block entailed the same 8 multiplication problems presented in random order. After each block participants were asked to approximate, on a scale from 0-100, how much they used retrieval or calculation strategies in that block. At the end of the first day, participants were asked to report on whether they think they received active or sham stimulation. Participants returned for a second day of testing, though these data are not analysed here, as they are beyond the scope of the present study.

Analysis was carried out on the accuracy and median reaction time for correct responses, and their report of the chosen strategy (retrieval vs. calculation).

### 2.4. tRNS

Participants were assigned to a stimulation condition using a variance minimisation algorithm based on their baseline arithmetic abilities^42^, which aims to allow better matching between groups in comparison to random allocation^43^. Thirty-eight participants received active high-frequency tRNS (100–500Hz) on the first day of the experiment (**Figure 2**). Stimulation was generated using a StarStim R32 (Neuroelectrics, Barcelona, Spain) and delivered via Ag/AgCl electrodes with 3.14cm^2^ surface area and carried to the scalp by conductive gel. tRNS was delivered over F3 and F4 according to the 10–20 system, with an amplitude of 1mA peak-to-peak for a duration of 20 minutes, beginning with the start of the arithmetic learning task. The tRNS had a ramp period at the beginning and end of the stimulation of 30 seconds. The remaining participants (n=37) received sham stimulation. Participants in the sham condition received 30 seconds of stimulation at the beginning of the task: 15 seconds of ramp up and 15 seconds of ramp down. To ensure experimenter blinding, the allocation software provided a pseudonymised stimulation group immediately following the baseline task, which were then matched with a blinding stimulation protocol in the stimulation software.

**Figure 2:**
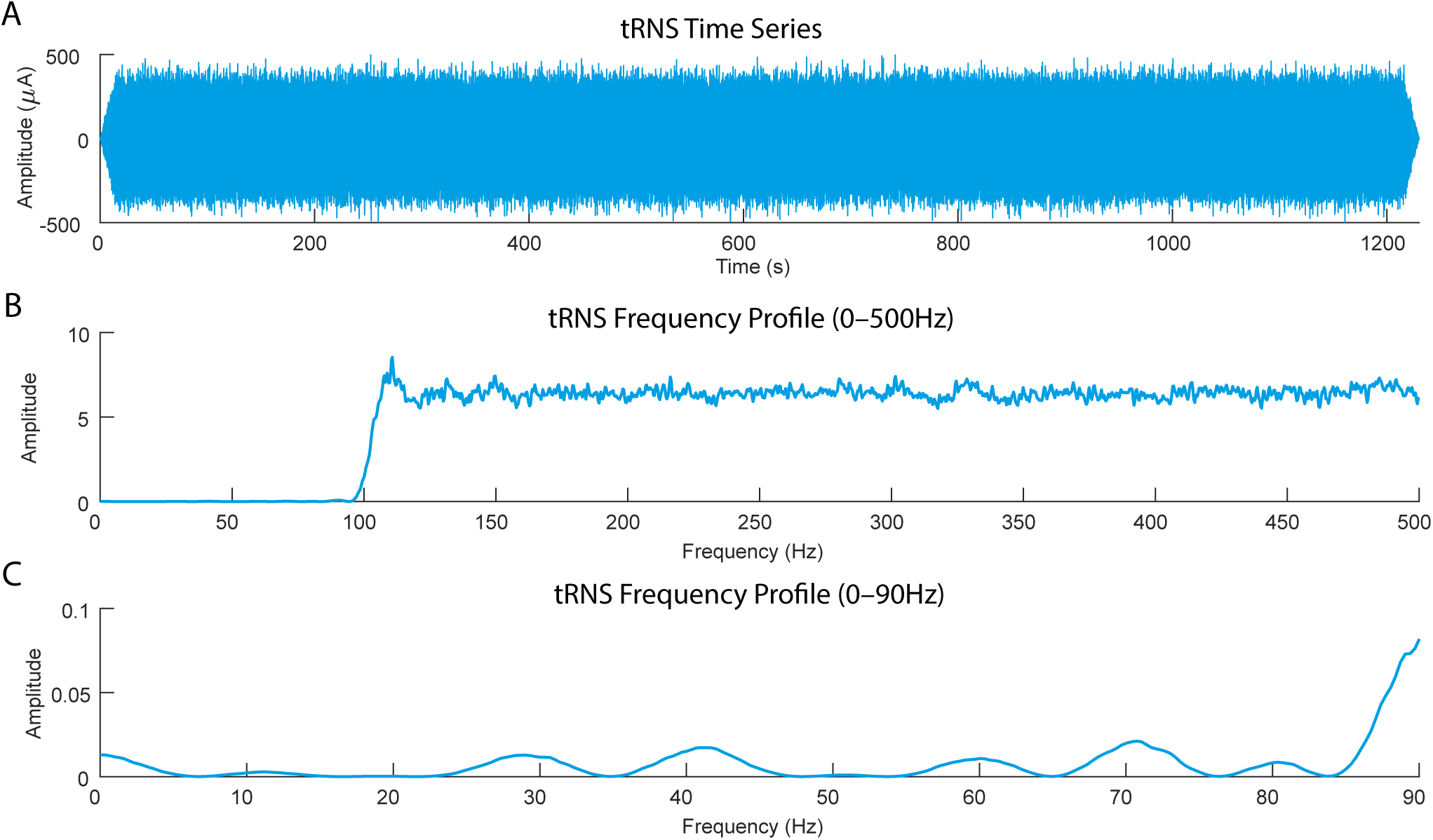
The profile of the tRNS stimulation. a) The time series of the stimulation over the 20-minute period, b) The full frequency profile (0–500Hz) of the tRNS stimulation, and c) the frequency profile in the 0–90Hz range. Despite the tRNS being filtered for higher frequencies (100–500Hz), peaks in the activity are visible in the lower frequency range (0–90Hz)

### 2.5. EEG collection

EEG data were collected via the stimulation unit from 28 sites whilst stimulation was ongoing and 30 sites once stimulation had ended (electrodes F3 and F4 were added) at a sampling rate of 500Hz. The data were recorded with Ag/AgCl electrodes. Similar to the stimulation preparation, EEG scalp sites were prepared by first abrading the scalp with abrasive gel, and then cleaned using surgical alcohol. The space between the scalp and electrodes was bridged using conductive gel.

### 2.6. EEG preprocessing

EEG data were imported into EEGLAB^44^ for preprocessing and were then filtered using a 0.5–80Hz band-pass filter. The mean was removed from each channel, and then periods of non-stereotyped artefactual data were removed after identification via visual inspection. Infomax ICA was run, and any periods of non-independence between components were identified by eye. Following removal of these periods of non-independence, we ran a second infomax ICA. For analysis, two versions of the EEG data were created. In the first version, we removed all components associated with stereotyped noise (e.g., blinks, lateral eye movements, heartbeat artifacts), as well as a component with a bilateral activation pattern and spectral profile matching the tRNS spectral profile. In the second version, only components associated with typical stereotyped EEG artefacts were removed whilst the tRNS artefact components remained, allowing us to investigate the impact of tRNS-induced artefacts alone.

For the ERP analysis, EEG data were grand averaged over trials. For the frequency analysis, the data were epoched into two-second windows with 50% overlap and a Hanning window was applied before being put through a FFT. To calculate the theta/beta ratio (TBR) for the data, theta (4-7.5Hz) and beta (15-30Hz) power was extracted relative to the average power in the 1.5-35Hz range. TBR was then calculated as the relative theta power divided by relative beta power. Intertrial phase coherence (ITPC) values were computed following methods outlined in Cohen (2014). Finally, to explore whether any TBR effects were better explained by changes in the underlying background noise (the aperiodic signal of the power spectrum; **Figure S1**), we extracted the exponent of the aperiodic signal. To extract the aperiodic signal from the resting and task EEG, the participants’ power spectra were exported from MATLAB and analysed using the FOOOF package (v 1.0.0)^46^ in Python (v3.7.0). The FOOOF package aims to breakdown the power spectrum into two components; the periodic signal, reflecting peaks in the spectrum defined by their frequency and power; and the aperiodic background signal, presenting as a 1/f power law and can be described by its offset (power at the lowest frequency) and its exponent, which is reported here. We computed the aperiodic signal over the 3-40Hz range, with a larger aperiodic exponent reflecting a greater slope in the background noise (relatively higher power in low frequencies than high frequencies).

### 2.7. Analysis

Behavioural analyses were conducted using linear mixed-effects models run via the *nlme* package^47^ and ordinal mixed-effects models were run using the *clmm* package^48^ in RStudio^49^. Plots for these models were created using sjPlot^50^.

RT data was analysed using linear mixed-effects models. Prior to data analysis, the median RT was log transformed (due to the skewness of the data) and z-scored to allow for calculation of standardised β coefficients. To ensure a linear relationship between block number in the task and RT, block number was also log transformed^51,52^. All other predictors were normalised to prevent issues with multicollinearity.

As accuracy data was calculated as a proportion of answers correct per block, with most participants getting 100% to 87.5% of answers correct per block (8 problems per block), data was levelled and heavily left skewed. Therefore, accuracy data was analysed using ordinal mixed-effects models.

Bayesian t-test^53^ using a Cauchy distribution with a scaling factor of δ-0.707 were used to compare the spectral profiles between the sham, tRNS removed, and non-removed data. As previously suggested^54^, Bayes factorss (BFs) above 3 were considered as moderate evidence (in favour of a model), values above 10 as strong evidence, and values above 100 as extremely strong evidence. EEG data were compared during both the online and offline periods. For both stimulation groups we defined the online period as the time in which the active tRNS group received stimulation, and the offline period as the time following tRNS until the end of the task. As an alternative method for analysis^55^, we conducted an analysis in which the data to inform the prior distributions to minimise impact of multiple comparisons. Instead of conducting Bayesian t-tests at each time point, peaks in the data are selected from the grand average, and then the prior distribution for the null is restrained by what is feasible for the dataset. The results from this analysis are reported in the **Supplementary Materials**.

## 3. Results

### 3.1. Stimulation blinding

Participants in the sham condition reported being in the stimulation condition at approximately the same rate as participants in the stimulation condition (78% vs 79%). We analysed the stimulation blinding data using Bayes factors as we were interested in providing evidence in favour of the null hypothesis, rather than only giving evidence to reject it as seen in frequentist analysis. The results from the two-tailed test demonstrated that the data were not favoured under one model over the other (BF_10_ =0.69), preventing us from drawing conclusions regarding the efficacy of our blinding.

### 3.2. Learning slopes

As specified in our preregistration to confirm that learning had taken place, we calculated the slope coefficient for block number on RT to identify participants with a non-negative slope (indicating no decrease in RT over the task period). In total, 5 participants demonstrated no decrease in RT, all from the sham group. A chi-squared test checking whether the two groups differed indicated that this was statistically significant (Χ^2^(1)=5.35, *p=*0.02). Originally, participants with a non-negative slope were to be excluded from the analysis. However, as this would lead to a large group imbalance between stimulation conditions, and would bias our results, these participants were left in.

### 3.3. RT

As there were no time constraints on participant response, participants were able to take as long as needed to ensure an accurate answer, though participants were encouraged to also respond quickly. In such cases, and as specified in the pre-registration, we would expect to observe the effect in the RT data rather than the accuracy data^16,56^. Results from the mixed effects model indicated that there was a significant 3-way interaction between the block number, stimulation group, and participants baseline RT (β=0.1. S.E.=0.03, t(1186)=2.16, p=0.03; **Figure 3, for full results see Table S2**). Dividing the participants in tercile groups based on the baseline RT and running a simpler model for each group (RT ~ block * stimulation group) revealed that participants with slower baseline RTs benefited from tRNS with faster RTs in earlier blocks (β=0.19, S.E.=0.06, t(395)=3.16, p=0.002). Though the sham and stimulation groups converged by the end of the experiment. This effect did not exist in the medium (β=−0.02, S.E.=0.04, t(394)=−0.39, p=0.69) or faster baseline RT groups (β=−0.03, S.E.=0.02, t(394)=−1.48, p=0.14). Note that there was no tRNS effect in baseline RT between active and sham stimulation in any of the tercile groups (fast baseline RT: β=0.08, S.E.=0.19, t(22)=0.43, p=0.66; medium baseline RT: β=−0.09, S.E.=0.13, t(22)=−0.74, p=0.46; slow baseline RT: β=−0.11, S.E.=0.15, t(22)=−0.73, p=0.47).

**Figure 3.**
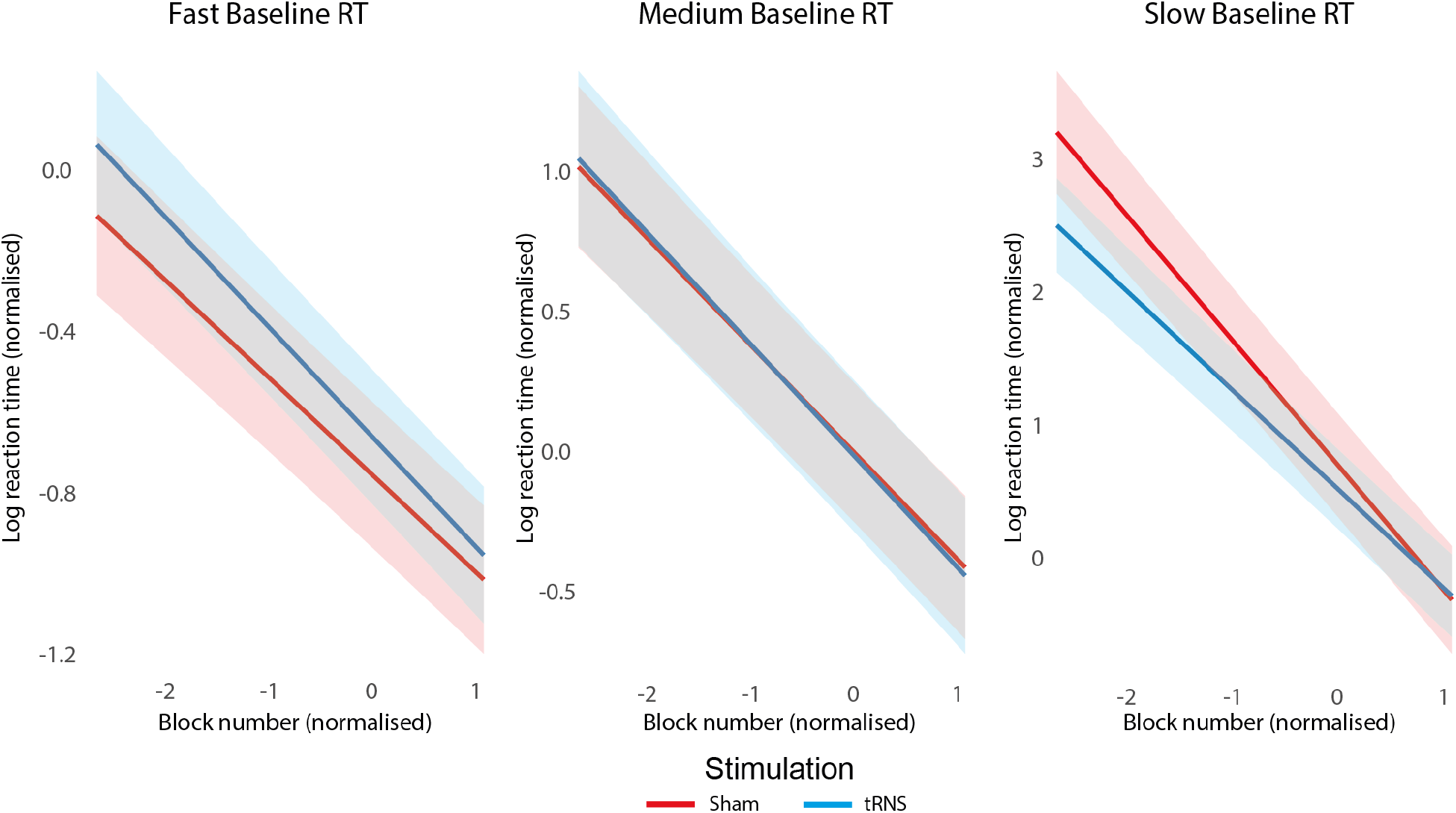
Predicted model fit results from the 3-way interaction between the block number, stimulation group, and participants baseline RT. Error bars represent 95% confidence intervals.

### 3.4. Accuracy

Results from the ordinal mixed effects model showed that there was a main effect of block (b=0.42, S.E.=0.09, z=4.42, p<0.001) and baseline accuracy (b=0.80, S.E.=0.17, z=4.70, p<0.001) on task accuracy. Accuracy increased over block and higher baseline accuracy was associated with higher task accuracy. However, there was no main or interacting effect of stimulation on participant accuracy (all p’s > 0.05; **for full results see Table S3**)

### 3.5. Calculation/retrieval results

Similar to the accuracy results, there was a main effect of block on the amount participants favoured retrieving the answers over calculating them (b=−0.62, S.E.=0.017, t(1186)=−35.5, p<0.001), with participants choosing to retrieve the answers more in later blocks. However, there were no main or interacting effects of participants’ baseline RT or stimulation group on the amount that the favoured either calculation or retrieval (all p’s > 0.05; **for full results see Table S4**).

### 3.6. tRNS artefact removal results

The results of the Bayesian t-tests indicated that whilst the grand average ERPs did not differ between the active and sham stimulation groups, regardless of whether we removed the tRNS artefact from the data or not, the standard deviations did. This problem was solved when we removed the tRNS artefact (BF_10_’s<3; **Figure 4**). Similarly, the frequency domain analysis of the data revealed substantial differences between the active and sham stimulation groups before tRNS scrubbing was applied. Specifically, tRNS was associated with several broad peaks (larger than the underlying power evoked in neural circuits) at approximately every 10Hz from 30–80Hz (aside from 50Hz). These peaks were not present in the sham data. Following ICA removal of the tRNS signal, the active tRNS data was largely comparable to the sham data but notably different from the non-removed data in both frontal and occipital sites (**Figure 5**).

**Figure 4:**
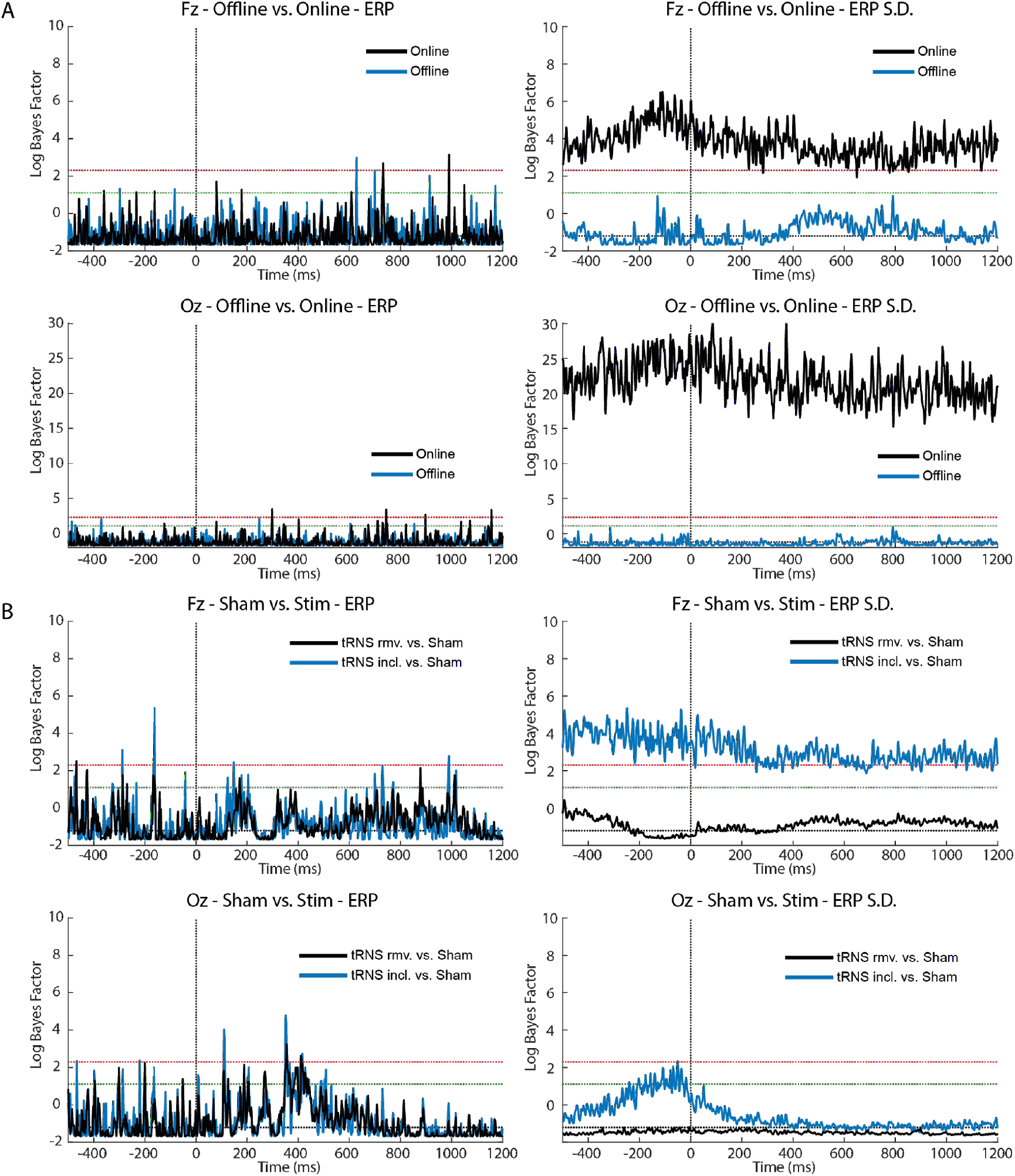
Time-domain results of Bayesian t-tests after tRNS artefact removal. A) Results of Bayesian t-tests (log BF_10_) comparing ERPs and ERP standard deviations (S.D.) for the data from the active tRNS group with the tRNS removed against the data with the tRNS artefact included. Results are for both electrodes Fz (closest to the tRNS electrodes over F3–F4) and Oz (furthest away from the tRNS electrodes over F3–F4). Red and green dotted lines represent a BF of log(3) and log(10), respectively, whereas the black dotted line represents a value of log(0.3). Note that the scale at Oz is larger than that of Fz. These results suggest that there were no differences in the ERP between online and offline periods regardless of whether the stimulation artefact was included or not, but that there was substantially more variance (S.D.) in the ERP when stimulation was ongoing. B) ERPs and ERP S.D. for the data sets with the tRNS artefact included (tRNS incl.) and tRNS artefact removed (tRNS rmv) compared against the sham participants for both Fz and Oz electrodes. As above, red, green, and black dotted lines represent a BF of log(3), log(10), and log(0.3), respectively. Similar to the online vs. offline comparison in panel A, there was little difference in the ERP between the sham and active tRNS conditions regardless of whether stimulation noise was removed or not. However, when the tRNS artefact was included, there was a noticeable increase in variance compared to the sham condition, but no increase in variance was noticeable when the artefact was removed.

**Figure 5:**
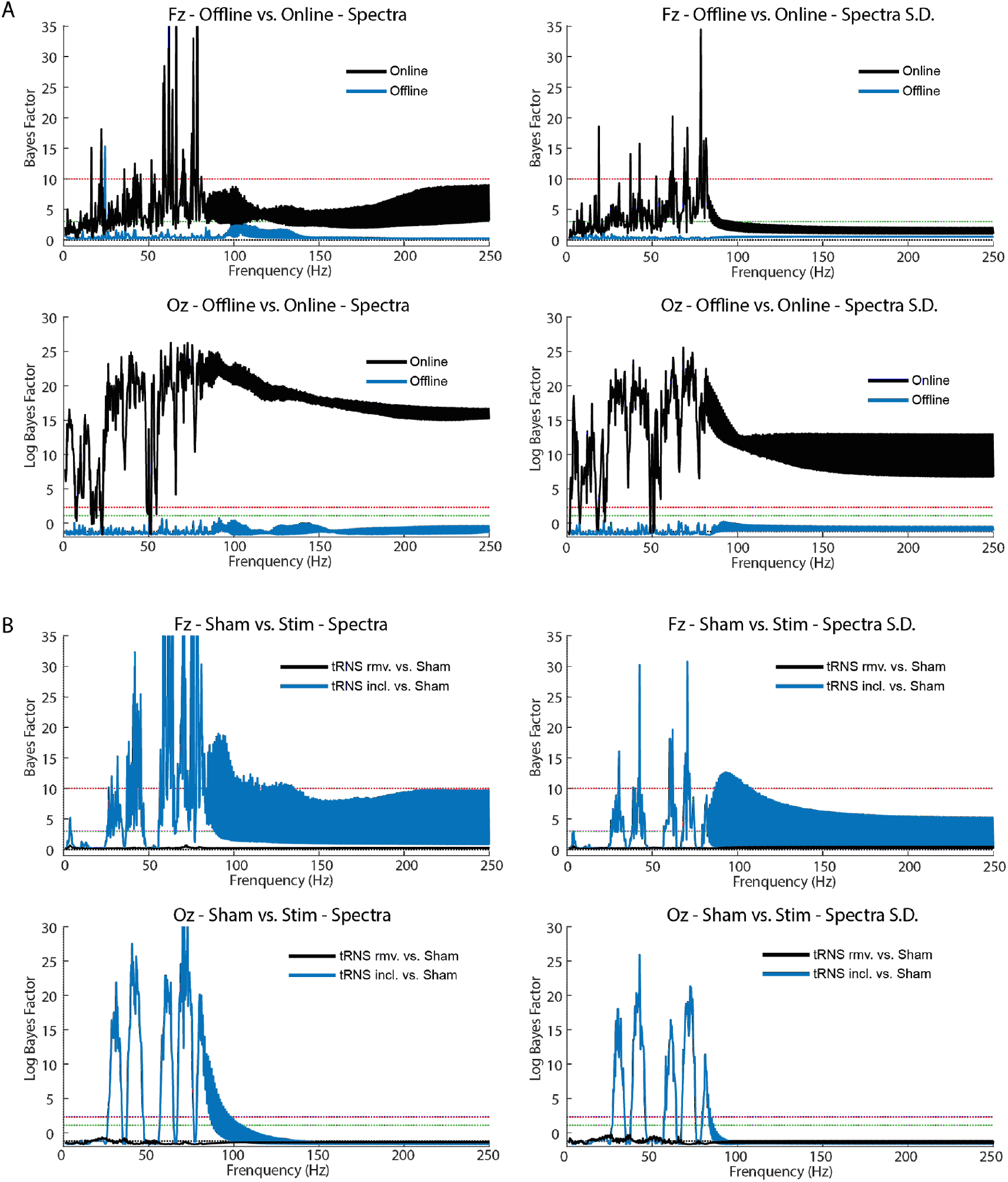
Frequency-domain results of Bayesian t-tests after tRNS artefact removal. A) Results of Bayesian t-tests (log BF_10_) comparing the spectral profile and their standard deviations (S.D.) for the data form the active tRNS group with the tRNS removed against the data with the tRNS artefact included. Results are for electrodes Fz (closest to the tRNS electrodes over F3–F4) and Oz (furthest midline electrode from the tRNS electrodes over F3–F4). Red and green dotted lines represent a BF of log(3) and log(10), respectively, whereas the black dotted line represents a value of log(0.3). Note that the scale at Fz is different from that of Oz, with the latter being a log scale. Unlike the ERP results, both the periodograms and their standard deviations were notably different between the data with the tRNS artefact included and the tRNS artefact removed during the online period but not during the offline period where the tRNS stimulation had finished. This is despite tRNS activity only being delivered in the 100–500Hz range. B) The spectra and their standard deviations for the data sets with the tRNS artefact included (tRNS incl.) and tRNS artefact removed (tRNS rmv.) against the sham participants for both Fz and Oz electrodes. As above, red, green, and black dotted lines represent a BF of log(3), log(10), and log(0.3), respectively. Note that the Oz results are reported on a log scale, whilst the Fz results are not. These results indicate that the presence of 100–500Hz tRNS noise led to substantial noise in the 0–100Hz range. However, this noise could be removed, leading to a spectral profile largely comparable to sham. Please note that the thickness in the lines in the top of row of both panel A and B from ~90Hz upwards represent compression of data points at varying BF values due to the wide variation in power across frequencies in the tRNS signal. This high compression of consecutive data points leads to visual artefact during plotting.

### 3.7. Theta/beta ratio

tRNS and sham groups did not differ in their resting TBR (b=0.22, S.E.=0.14, t=1.64, *p*=0.1). We did not observe significant main effects of stimulation (b=0.63, S.E.=0.36, t=1.71, p=0.09) or an interaction between stimulation and baseline TBR interaction (b=−0.29, S.E.=0.21, t=−1.35, p=0.18) when predicting participants’ online TBR.

#### RT

There was a significant interaction between stimulation group and TBR recorded during both the pre-task resting period (β=−0.21, S.E.=0.1, t(59)=−2.06, p=0.04) and the online period (β=−0.25, S.E.=0.1, t(56)=−2.46, p=0.02). In both cases, active stimulation was associated with faster RTs amongst participants with higher TBRs (**Figure 6a**). Post-hoc analysis of the simple slopes revealed high TBR lead to reduced reaction times when the participant received active stimulation (b=−0.16, S.E.=0.06, t(59)=−2.56,p=0.01), but not when they received sham stimulation (b=0.04, S.E.=0.07, t(59)=0.59, p=0.55). For full results see **Table S5 & S6**.

**Figure 6.**
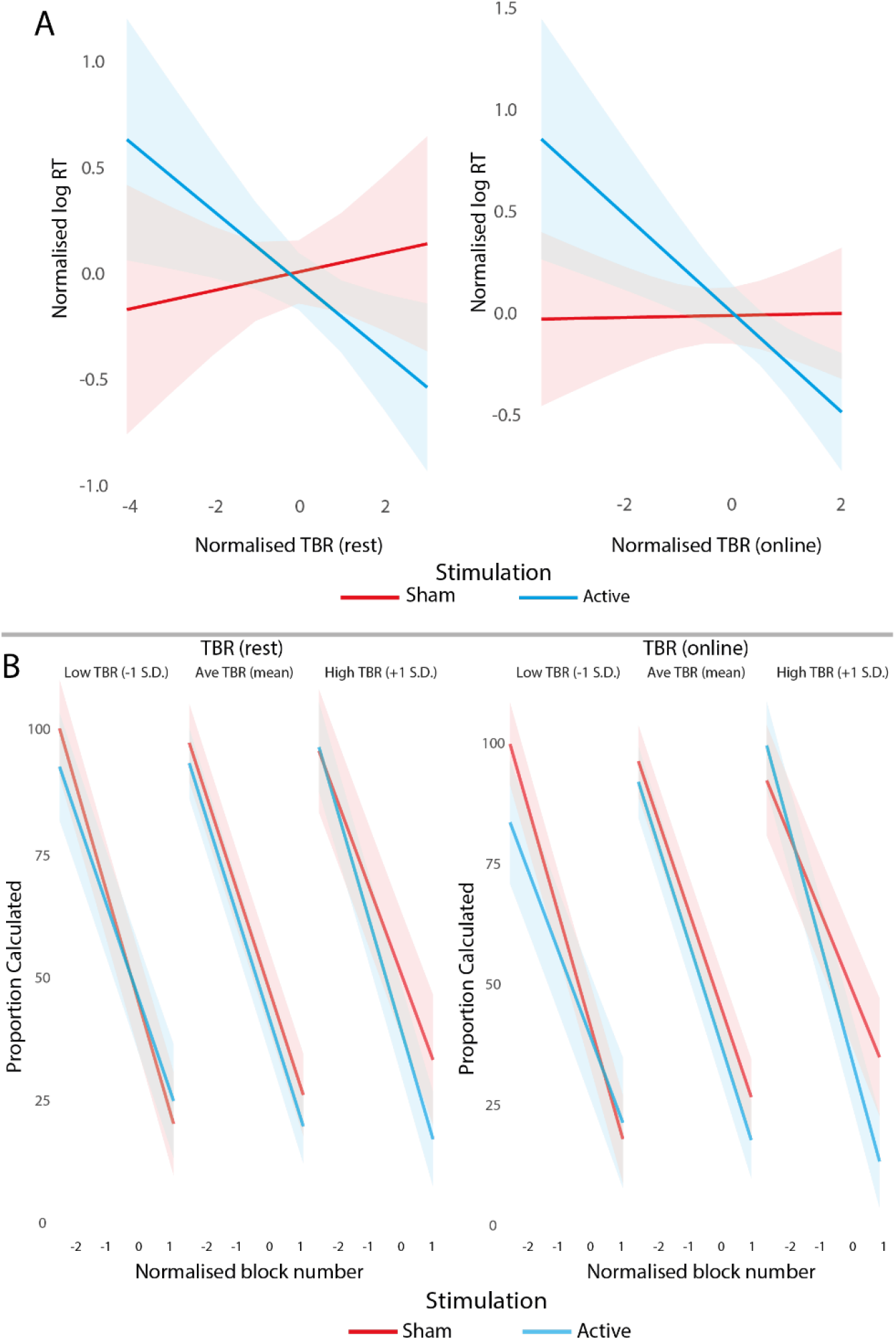
The effects of stimulation and TBR on behavioural outcomes. A) Relationship between median RT and TBR during the resting period (left) and during stimulation (right). In both cases, higher TBR were associated with better RTs in the active stimulation group, but not in the sham group. B) Proportion calculated as a product of block number, stimulation, and TBR. During rest (left) and the online stimulation period (right), for participants with high TBR active stimulation led to greater levels of retrieval in later blocks compared to sham stimulation. Moreover, for participants with low TBR values during the online period, participants receiving active stimulation were more likely to report retrieving results than their sham counterparts.

#### Accuracy

Results from the accuracy models indicated that the 3-way interaction between block number, stimulation group, and online TBR was significant (b=−0.33, S.E.=0.15, z=−2.28, p=0.02, **Figure 7**), but did not reach significance when using the 3-way interaction with resting state TBR (b=−0.08, S.E.=0.13, z=−0.62, p=0.53). tRNS was less beneficial for those with low TBR individuals. These individuals were less likely to answer correctly at the beginning of the training (approximately the first half of the blocks). Later, these participants’ accuracy increased to 100% correct answers. By comparison, their sham counterparts were more likely to get 100% of answers correct during the early stages of the task. In contrast, tRNS was beneficial to high TBR participants who were more likely to answer 100% of problems correctly throughout the experiment, whereas their sham counterparts were less likely to get 100% in the earlier blocks (~75% vs. ~60%). For full results see **Table S7 & S8**

**Figure 7.**
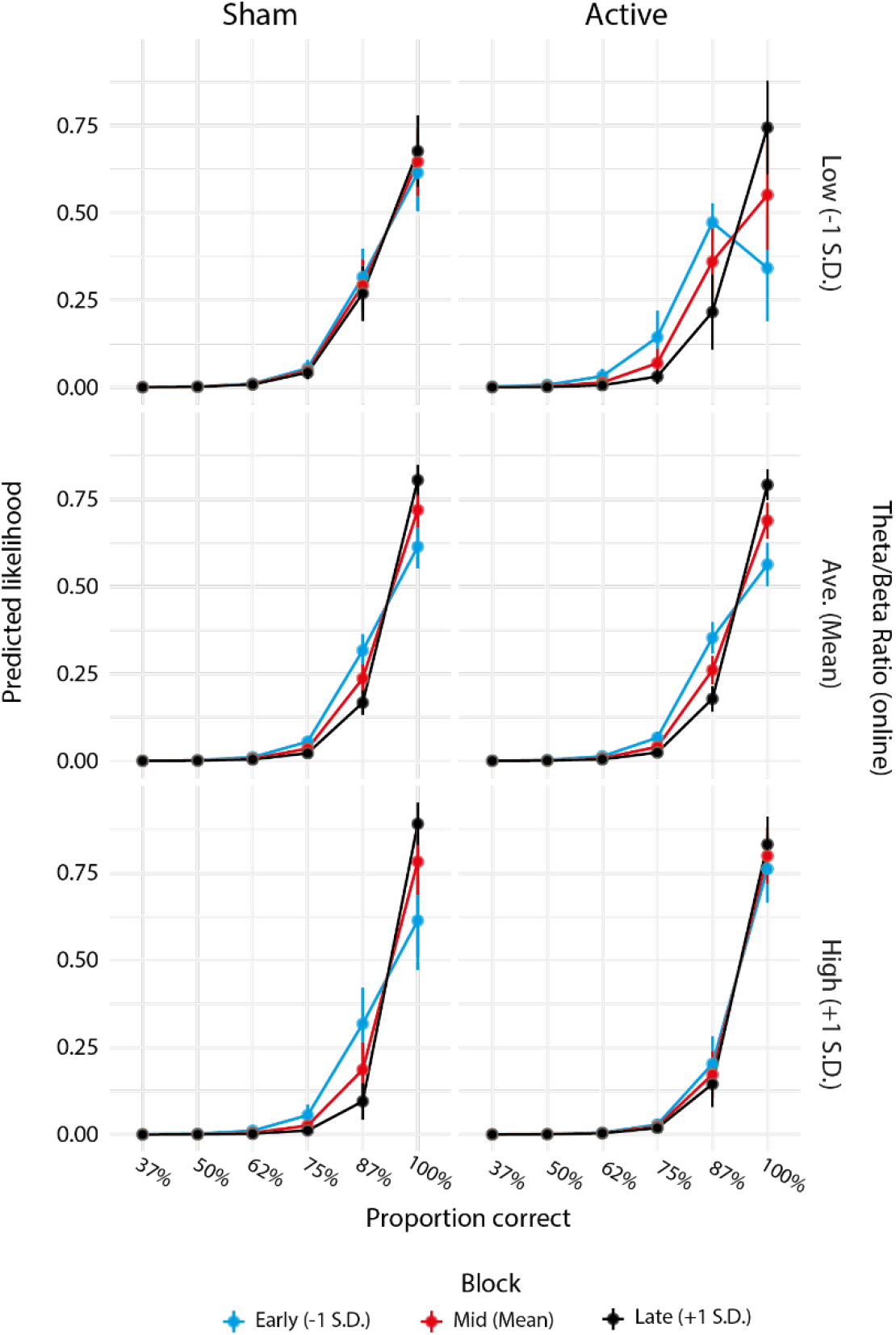
The ordinal mixed effects model results regressing accuracy back on block number, stimulation group, and online TBR. Each graph shows the predicted likelihood of participants getting that proportion of responses correct in a block. For participants with low TBR (top right plot) active stimulation was associated with lower levels of accuracy in earlier blocks compared to sham stimulation, though accuracy improved to match sham by later blocks. To a lesser extent, participants with high TBR benefited from active stimulation demonstrating slightly higher accuracy in the earlier blocks than those who received sham stimulation

#### Calculation/Retrieval

The proportion of answers calculated vs. retrieved was impacted by the 3-way interaction between block number, stimulation group, and TBR during both the resting state period (b=−3.96, S.E.=0.79, t(1084)=−4.96, p<0.001) and during stimulation (b=−6.53, S.E.=0.83, t(1032)=−7.81, p<0.001; **Figure 6B**). As in the case of accuracy and RT, tRNS was beneficial for those with high resting TBR in the later blocks leading to a greater proportion of answers retrieved compared to sham (b=−15.58, S.E.=7.63, t(59)=−2.04, p=0.04), which was not seen in the early (b=−6.43, S.E.=7.63, t(59)=−0.84, p=0.40) or mid blocks (b=−11.01, S.E.=7.55, t(59)=−1.45, p=0.15).A similar effect was noted for participants with a high online TBR, who benefited from active stimulation over sham in later blocks (b=−21.09, S.E.=7.18, t(56)=−2.93, p=0.004), showed a trend in mid blocks (b=−13.30, S.E.=7.09, t(56)=−1.87, p=0.06), but not in early blocks (b=−5.52, S.E.=7.18, t(56)=−0.76, p=0.44). For full results see **Table S9 & S10**.

### 3.8. Aperiodic signal analysis

The relationship between TBR and behavioural outputs may instead be explained in differenced in the underlying aperiodic signal rather than theta and beta band specific changes^29,57^. To explore this, we extracted and analysed the aperiodic exponent from the resting and online power spectra. The exponents were then included as predictors in the mixed-effects models predicting RT, accuracy, and calculation vs. retrieval as in the case of TBR.

#### RT

There was a 3-way interaction between block number, stimulation group, and participants resting aperiodic exponent (β=−0.12, S.E.=0.02, t(1084)=−5.09, p<0.001; **Figure 8A**). For participants with a high resting state aperiodic exponent, which is suggested to reflected low E/I ratio^32^, active stimulation was most effective leading to faster RTs compared to sham stimulation as the number of block increased (early blocks: b=−0.17, S.E.=0.14, t(59)=−1.21, p=0.23, mid blocks: b=−0.33, S.E.=0.13, t(59)=−2.39, p=0.02, late blocks: b=−0.49, S.E.=0.14, t(59)=−3.45, p=0.001). In contrast, for participants with a low resting state aperiodic exponent, active stimulation lead to slower RTs compared to sham stimulation as the number of block increased (early blocks: b=0.17, S.E.=0.13, t(59)=1.24, p=0.22, mid blocks: b=0.24, S.E.=0.13, t(59)=1.81, p=0.07, late blocks: b=0.31, S.E.=0.13, t(59)=2.28, p=0.02).

**Figure 8.**
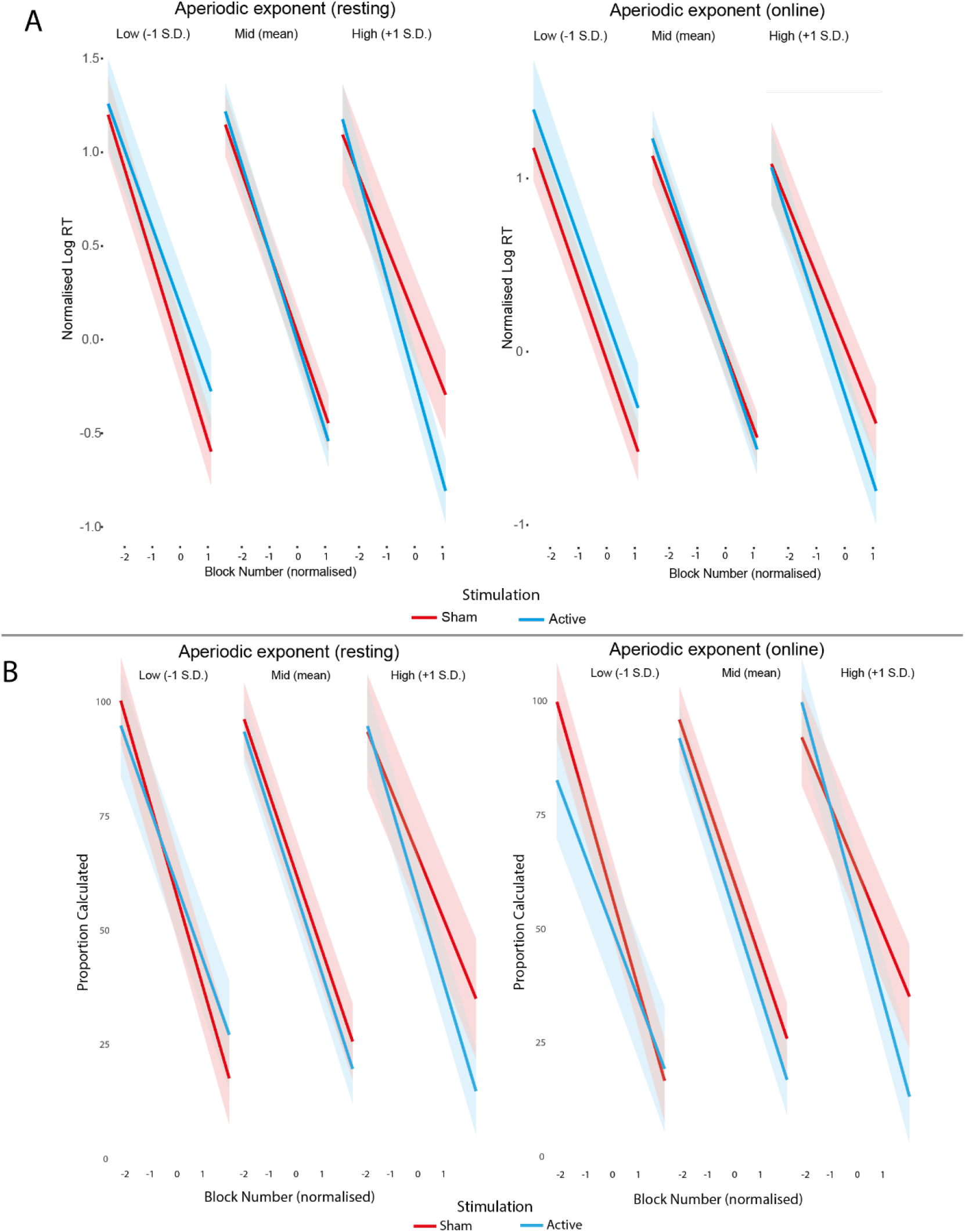
Figure demonstrating the relationship between aperiodic exponent, both at rest and during stimulation, and stimulation on behavioural outcomes. A) The relationship between aperiodic exponent, block number and stimulation on RT. Participants with high aperiodic exponent benefitted most from active stimulation, leading to faster RTs by the end of the experiment. By comparison, low aperiodic exponent individuals did not benefit from either stimulation condition. B) Similar to the RT data, proportion retrieved was influenced by the 3-way interaction between stimulation, aperiodic exponent, and block number. Specifically, participants with high aperiodic exponent benefited from active stimulation, leading to a higher proportion of answers retrieved in later blocks compared to the sham stimulation counterparts.

The 3-way interaction was also significant for the online aperiodic exponent (β=−0.05, S.E.=0.02, t(1033)=−2.17, p=0.03). Once again, compared to sham stimulation, active stimulation was more beneficial for participants with high online aperiodic exponent as the number of block increased (early blocks: b=−0.17, S.E.=0.13, t(56)=−1.28, p=0.2, mid blocks: b=−0.27, S.E.=0.13, t(56)=−2.06, p=0.04, late blocks: b=−0.37, S.E.=0.13, t(56)=−2.71, p=0.009). The opposite pattern was found for those with a low online aperiodic exponent, showing an overall trend towards slower RTs for active vs. sham stimulation in mid and later blocks (early blocks: b=0.23, S.E.=0.14, t(56)=1.57, p=0.12, mid blocks: b=0.24, S.E.=0.14, t(56)=1.68, p=0.09, late blocks: b=0.25, S.E.=0.14, t(56)=1.69, p=0.09). For full results see **Table S11 & S12.**

#### Accuracy

The only significant interactions that included stimulation as a predictor was between stimulation group and aperiodic exponent (b=0.69, S.E.=0.33, z=2.12, p=0.03; **Figure 9**). Stimulation was associated with a greater proportion of correct answers for participants with a high aperiodic exponent compared to those with a lower aperiodic exponent. For full results see **Table S13 & S14.**

**Figure 9.**
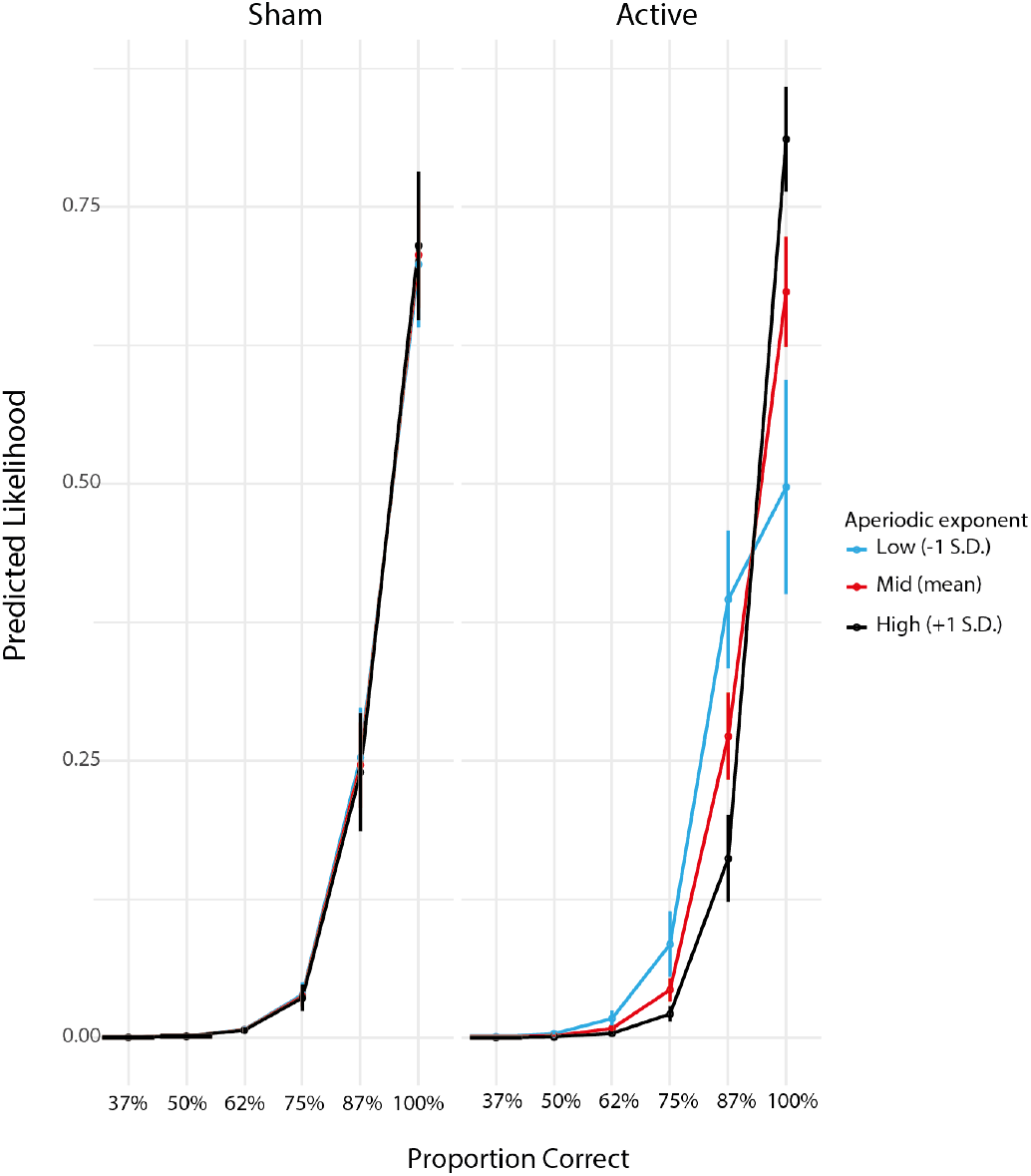
Results from the stimulation by aperiodic exponent interaction on accuracy, where the effect was primarily seen in the active stimulation condition. Individuals with high aperiodic exponent demonstrated higher accuracy than their low aperiodic exponent counterparts when they receive active stimulation. In the sham condition, high and low exponent participants were similar in performance.

#### Calculation/Retrieval

There was a significant 3-way interaction between block number, stimulation group, and the resting aperiodic exponent (b=−4.92, S.E.=0.79, t(1084)=−6.20, p<0.001; **Figure 8B**). Analysis of the simple slopes indicated that this effect was primarily in the participant with a high aperiodic exponent. In the mid (b=−13.71, S.E.=7.46, t(59)=−1.83, p=0.07) and late (b=−19.52, S.E.=7.55, t(59)=−2.58, p=0.01) blocks, active stimulation was associated with greater proportion retrieved than those with a high exponent who received sham stimulation. However, this was not the case in the early blocks (b=−7.91, S.E.=7.55, t(59)=−1.04, p=0.29). The online aperiodic exponent also interacted with block number and group label to predict the proportion of answers participants retrieved (b=−6.67, S.E.=0.85, t(1033)=−7.88, p<0.001). Once again, simple slopes analysis indicated that this effect was in the participant with a high aperiodic exponent, where active stimulation was associated with greater proportion retrieved than those who received sham stimulation in mid (b=−12.14, S.E.=6.99, t(56)=−1.73, p=0.08) and late (b=−20.14, S.E.=7.08, t(56)=−2.84, p=0.006) blocks. However, this was not the case in the early blocks (b=−4.14, S.E.=7.08, t(56)=−0.58, p=0.56). For full results see **Table S15 & S16**

Notably, when we included the TBR or the aperiodic exponent as a covariate in the models that reported effects for TBR (aperiodic exponent as a covariate) or aperiodic exponent (TBR as a covariate) in predicting behavioural outcomes, the reported effects remained significant, therefore highlighting their independent moderating effect.

### 3.9. ERP results

Following the successful removal of the tRNS artefact, ERP plots of the tRNS-removed data from two midline sites revealed two main differences between the active and sham tRNS groups, defined as time periods in which the Bayes factor of the group difference was greater than 3. First, over frontal sites, an early negative potential, peaking 100ms following stimulus presentation, was greater amongst the active tRNS group compared to the sham group. Notably, this difference was not seen following the presentation of a fixation cross (**Figure 10**). This indicates that the effect is specific to stimulus presentation, rather than a general tRNS-induced artefact, and highlights our success in removing the tRNS-induced artefact.

**Figure 10:**
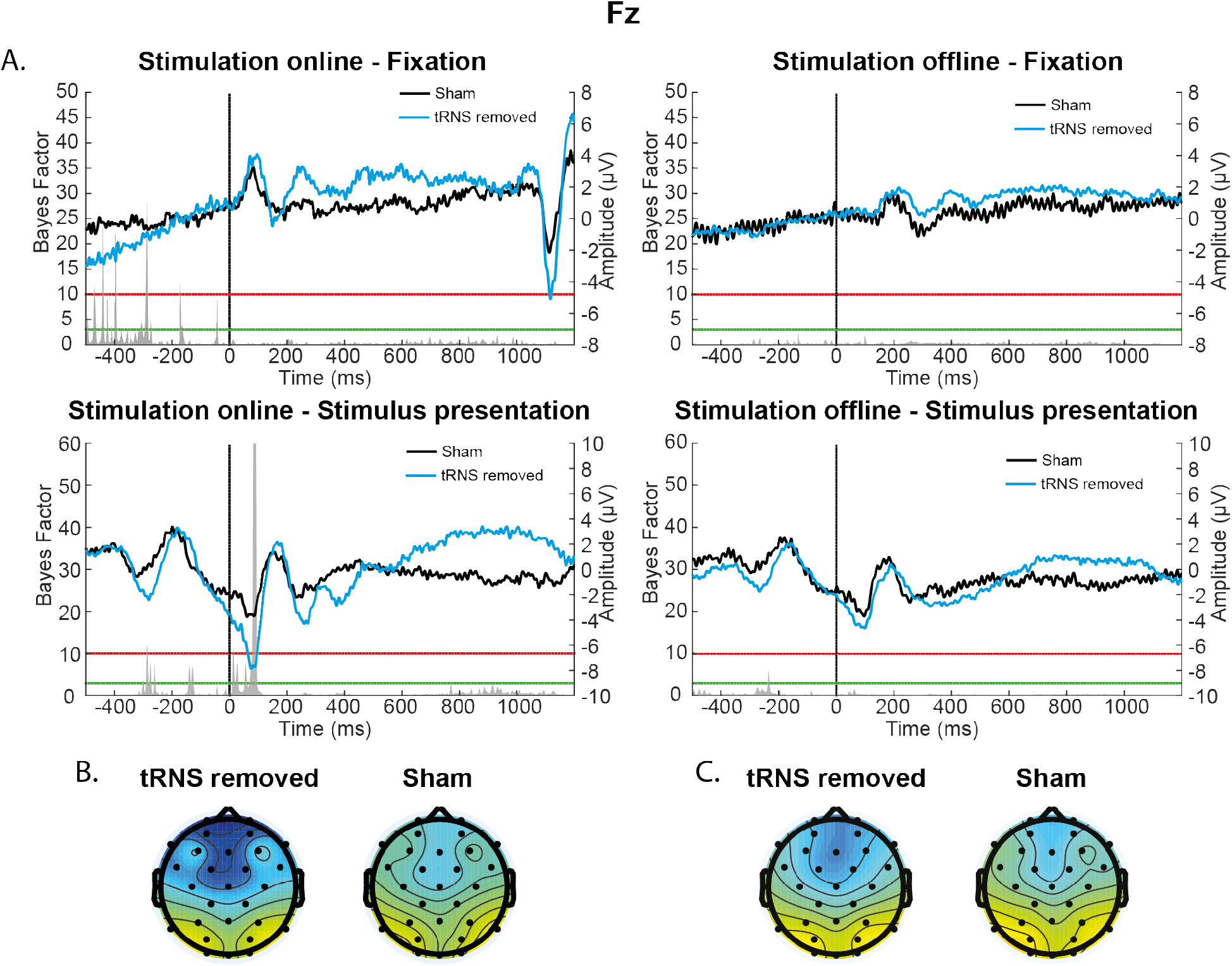
Task-related ERP effects at Fz following stimulus presentation. A) ERPs following stimulus and fixation cross presentations showing a stimulation group effect during the online, but not the offline, period following stimulus presentation. Underneath the ERP trace plots, the BF comparisons of the tRNS removed against the sham group. They highlight differences in the ERP waveform between the tRNS and sham groups following stimulus presentation, primarily in early ERPs (~100ms following stimulus presentation). Topographic distribution of activity (in μV) for the sham and tRNS artefact removed data at the 100ms time point during the online (B) and offline period (C). The colour bar range for all topographies is the same.

Second, at Oz, a greater positive inflection in the waveform 200ms after stimulus presentation was seen in the active tRNS group compared to the sham group (**Figure 11**). In contrast to the N1 effect, the group difference over occipital sites 200ms after stimulus presentation shows a lasting effect, as reflected in a similar effect during the offline period. During the offline period, the active tRNS group demonstrated greater positivity over occipital sites beginning approximately 200ms after stimulus presentation and continuing for another 300ms, likely reflecting the late positive potential (LPP). However, in contrast to all the other ERP results, this offline effect did not remain following multiple comparison correction (see the Supplementary Results section for a detailed description of the ERP multiple comparisons results).

**Figure 11:**
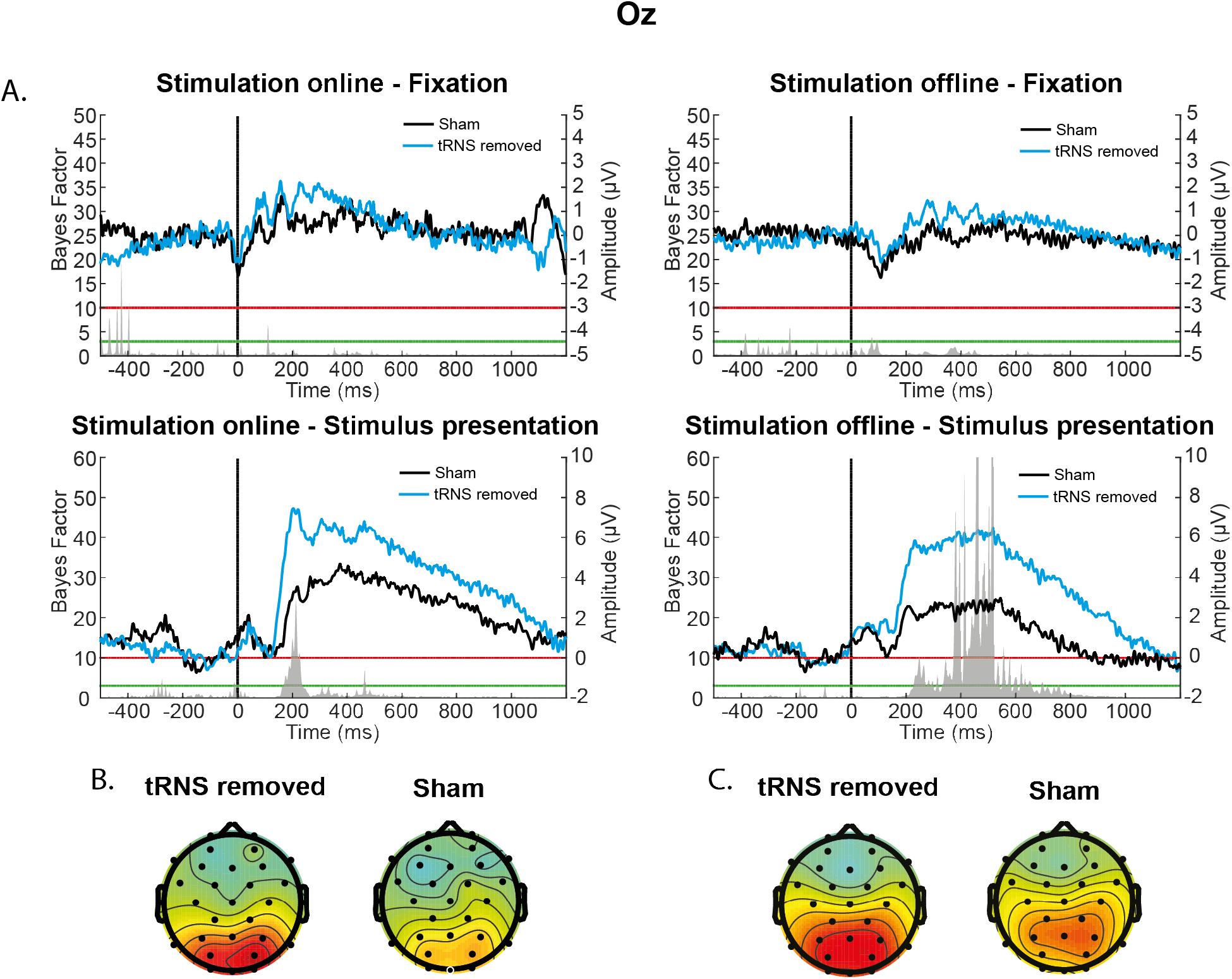
Task-related ERP effects at Oz following stimulus presentation. a) ERPs following stimulus and fixation cross presentations during both the online and offline periods. Waveforms are plotted for the tRNS artefact removed data and sham. Underneath the ERP trace plots, the BF comparisons of removed tRNS artefact data against the sham group are shown. They highlight differences in the ERP waveform between the tRNS and sham groups following stimulus presentation in both early ERPs (~200ms following stimulus presentation) and late ERPs (~400ms following stimulus presentation). B) Topographic distribution of activity (in μV) for the sham and tRNS artefact removed data at the 200ms time point during the online (B) and offline period (C). The colour bar range for all topographies is the same.

At both sites, ERP differences existed between the active and sham tRNS groups regardless of whether tRNS scrubbing had taken place, though this was less clear in the topographical maps due to stimulation noise (to see the tRNS removed, non-removed, and sham data plotted together, see **Supplementary Materials Figure S2 and S3**).

## 4. Discussion

Given the beneficial effects of tRNS on perceptual and cognitive performance, including cognitive training, we examined how tRNS may alter and interact with neural activity while participants completed a mathematical learning task and undergoing concurrent tRNS-EEG. During analysis, we removed the tRNS artefact from the ongoing EEG data and examined the underlying EEG signal change due to tRNS, and specific neurophysiological moderators, namely, TBR and aperiodic signal.

At the behavioural level, we found that tRNS influenced performance by impacting learning slopes and RTs. The RT effect was qualified by individual differences, as seen in recent tRNS studies. Notably, we found that participants with the slowest baseline RT benefitted most from stimulation, with improvements in performance occurring almost immediately with stimulation. By comparison, those with fastest baseline RT did not benefit from stimulation. Overall, these findings indicating that the beneficial effects of tRNS are dependent on individual differences at the behavioural level.

One of the novel contributions of our study is the effective removal of the tRNS artefact from the concurrently recorded EEG activity in order to provide an understanding of the neural mechanisms underlying the effect of tRNS on behaviour. When looking at the time-domain data, this procedure successfully reduced the variance of the ERPs in the case of tRNS, which was considerably larger in this group due to the tRNS artefact. Similarly, the tRNS artefact removal appeared successful in the frequency domain, leading to a periodogram similar to that seen in the sham group. This indicates that our procedure can be used to clean up both the time and frequency domains, which allowed us the sensitivity to detect neurophysiological effects and foster mechanistic understanding.

In line with the prediction in our preregistered analysis, and consistent with the same moderating effect of TBR on tRNS in sustained attention task^24^, we found that TBR over frontal sites moderates the effect of tRNS. Namely, tRNS was most beneficial for participants with high TBR. This effect was characterised by quicker RTs, higher accuracy during the early blocks, and faster shift to reliance on retrieval strategies compared to calculation, indicating greater proficiency when compared against their sham group counterparts. In contrast, low TBR participants who received tRNS had slower RT and poorer accuracy during the early blocks of the task, which might be due to the use of premature retrieval strategies early on compared to the sham participants. In contrast to the beneficial effect between active and sham tRNS in those with high TBR, the detrimental effect of active compared to sham tRNS on those with low TBR disappeared towards the end of the training. Given that low TBR is thought to reflect the successful top-down regulation of the dorsolateral prefrontal cortex^25^, participants with low TBR are likely to perform better without intervention as they are already working close to their optimum. In this case, tRNS seems to induce a transient impairment by shifting the reliance on an inadequate strategy (rote retrieval) that is not beneficial at this stage. In a similar study using the same task, we replicated the 3-way interaction between block, stimulation type, and baseline RT, as well as the effect of TBR and stimulation type on behavioural performance^58^.

Whilst TBR is assumed to reflect imbalance between theta and beta activity, it could be better explained by changes in the background noise of the brain, reflected by the aperiodic signal of the power spectrum. When we replaced TBR in the models with the aperiodic exponent, we found that the aperiodic exponent interacted with stimulation condition to predict RT, accuracy, and the participants’ reported retrieval in much the same way as TBR. Across these outcome measures tRNS benefitted participants with a higher aperiodic exponent. High aperiodic exponents are thought to reflect lower E/I ratios^30,31^. However, the effects of TBR and aperiodic exponent on any behavioural measure remained whilst controlling for the other, suggesting that they are capturing different elements of the underlying neurophysiological process.

Given the suggestion that tRNS impacts neuronal excitability^14,32–34^, the beneficial effects of tRNS on participant performance being dependent on participants’ natural E/I ratio could explain the mixed efficacy of tRNS in the previous literature. The balance in E/I ratio is what dictates the criticality of neural systems^59,60^, which is thought to be important for organising neuronal processing across frequency bands as well as generating rhythmic activity^61^. Amongst participants with more optimal E/I ratio at rest, such as those with a lower aperiodic exponent, the application of tRNS should throw off the E/I balance causing hypercriticality in the brain, a disorganised neural state that at the extreme end is linked with epileptic seizures^35,62,63^. By comparison, participants with a lower E/I ratio will benefit from tRNS as it should bring them closer to critical state. Moreover, there is evidence to suggest that the E/I ratio of a system may impact the power of high frequency activity^60,64^. Stimulation applied over frontal areas may lead to increases in subpopulations of neurons firing at a higher frequency. For participants with a high TBR, this may be beneficial as it reduces the relative number of neuronal populations firing at low frequencies, leading to a lower TBR, and greater cognitive control. However, amongst low TBR populations, this leads to greater imbalance as power in higher frequency bands is increased.

Broadly, the behavioural and neurophysiological results are in-line with stochastic resonance. At the behavioural level, tRNS may be improving the performance in participants with lower cognitive ability, as their poorer performance is assumed to be characterised by lower signal-to-noise ratio. By comparison, tRNS is adding noise in participants with more optimal signal-to-noise ratio, as reflected by high baseline performance, leading to little, if any, benefit of stimulation. However, our results only provide evidence for half of the predicted U-shaped function; participants with a lower signal-to-noise ratio improved in the presence of noise. Stronger evidence would be provided via mapping of the other half of the U-shaped function by demonstrating that a group of even lower signal-to-noise ratio participants do not benefit from stimulation at all. That said, the present effect could be explained by other mechanisms rather than stochastic resonance^3^. Future studies could test multiple tRNS amplitudes to better understand how behavioural response changes as a function of an interaction between tRNS amplitude and aperiodic exponent or baseline cognitive ability, to test more directly whether tRNS effect is due to stochastic resonance.

We also found notable differences in the time-domain EEG signal following stimulus presentation between the active and sham tRNS groups, which depended on whether the stimulus was presented during the online or offline period. The dissociation between the online and offline periods highlights the importance of removing tRNS artefacts in order to examine their immediate, rather than delayed, effect. During the online period, we found that active tRNS yielded a greater negative potential over frontal sites approximately 100ms following stimulus presentation than the sham condition. This effect was specific to the time of presenting the arithmetic equation and was not seen following fixation cross presentation. The N1 has been considered an attentional component reflecting perceptual attentional processes^65^, with higher attentional focus amongst participants with greater N1 amplitudes. In our sample, the increased N1 amplitude amongst tRNS participants following stimulus presentation may indicate greater attentional resources devoted to the problem presented. At a later time-point, over occipital sites, a larger P2 was observed amongst tRNS participants compared to sham participants. Again, this effect was specific to the stimulus presentation, and was more robust during the online period, as the P2 effect during the offline effect did not remain following multiple comparison correction. Similar to N1, P2 has often been associated with attentional processes^65^, with previous work identifying a parietal N1-P2 complex associated with early perceptual attention gating^66^. Moreover, the P2 was associated with increased theta phase locking over parietal sites, which was suggested to reflect attentional and working memory processes^67^. However, whilst applying tRNS resulted in increased P2 amplitude in the tRNS group compared to the sham group, this alteration in P2 was not associated with similar differences in the theta-band ITPC (see **Supplementary Materials**).

Overall, the TBR, N1, and P2 effects converge to the conclusion that the effect of tRNS when applied above the dlPFC is characterised by EEG markers that are associated with greater attentional resources. Given that we matched our participants based on their baseline arithmetic performance and that active tRNS yielded better performance than sham, a likely interpretation is that improvement in performance caused by tRNS reflects greater allocation of attentional resources during the training. Such mechanistic explanation clarify the relative broad improvement seen by tRNS in different tasks, including its effect on learning, where attentional resources play a key role^6^, as well as its potential benefit in attention-deficit/hyperactivity disorder^68^.

While we should be cautious with interpretations that stimulation is affecting the attentional network, as this could be a case of reverse inference^69^, given the experimental context, interpreting the results in relation to attention is not unreasonable^70^. Learning depends heavily on attention^71–73^, and as such, we might expect that modifying attention, in this case using tRNS, could benefit learning. In addition, the involvement of resting state TBR, which we acquired before the application of tRNS, further supports an attention-related interpretation. Indeed, the association between resting state TBR and baseline arithmetic performance was unreliable (r=0.04, p=0.72), excluding its role in arithmetic performance per se. Furthermore, within the context of the broader literature, stimulation changes in general rather than task-specific processes are in line with the effect of brain stimulation above the dlPFC on different cognitive tasks^5^. Thus, based on the electrophysiological results and the broader literature, the conclusion that stimulation led to early attentional changes provide the most parsimonious explanation for our results,

Finally, we also saw greater slow wave components, likely reflecting the LPP, in the active tRNS over occipital sites compared to the sham tRNS. This effect, compared to the earlier ERP components, was only observed during the offline period. LPP has been previously identified in memory recognition experiments and is thought to be reflective of conscious recollection^74,75^. For example, larger amplitudes have been observed when participants were presented with information they knew compared to unfamiliar information^76^. Whilst LPP is less well investigated in the mathematical learning and mental arithmetic literature, studies from other domains have highlighted its modulation by top-down control processes and have suggested that it originates from a bidirectional prefrontal-occipitoparietal co-dependency^77^. This is reflective of the mathematical learning literature, which has suggested that the movement from calculation to retrieval is associated with changes in neural activation from frontal to parietal sites^78–81^. However, whilst this interpretation could explain the faster RT amongst the tRNS participants, it does not correspond to the participants’ self-reported retrieval when the effect of stimulation was found only when TBR or aperiodic exponent were included.

By introducing a procedure to remove the artefact caused by tRNS we provide a novel understanding about how tRNS improve cognitive performance. An increased understanding of the mechanisms involved in brain stimulation, and its behavioural and neurophysiological moderating factors at rest and during stimulation would allow more effective stimulation protocols to improve cognition.

## 5. Supplementary Materials

### Supplementary Methods

**Table S1:** List of potential operands that can be combined to create the arithmetic problems presented to the participants.

| First<br>Operand |
| --- |
| 29 |
| 23 |
| 13 |
| 19 |
| 28 |
| 29 |
| 17 |
| 26 |
| Second<br>Operand |
| 3 |
| 4 |
| 2 |
| 6 |
| 8 |
| 7 |

**Figure S1.**
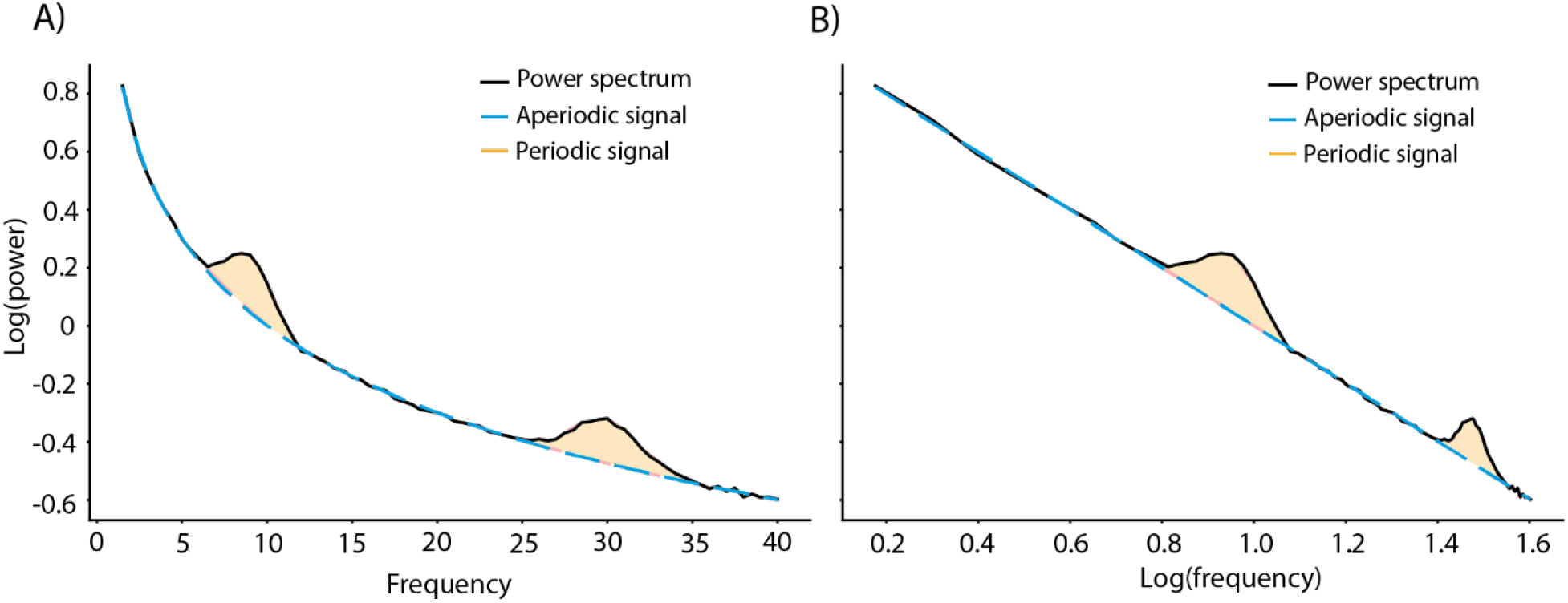
An example of how the FOOOF package breaks down the power spectrum (black line) into the periodic and aperiodic signals. A) The rhythmic oscillations (the periodic signal) plotted at 8 and 30Hz (yellow areas) are examples of EEG power signal typically analysed. The aperiodic signal (the line from which the peaks originate, here represented with dashed blue line) reflects the background noise of the brain and can be characterised by an exponent value. B) The aperiodic signal acts as a power law and therefore it can be plotted linearly in log-log space

### Supplementary Results

#### Behavioural results

**Table S2.** Results from the mixed-effects model regressing RT back on block number, stimulation group, and the baseline RT.

| Predictor | $\beta$ | S.E. | df | t | p |
| --- | --- | --- | --- | --- | --- |
| Intercept | 0.00 | 0.07 | 66 | 0.03 | 0.97 |
| **Block** | **-0.50** | **0.02** | **1186** | **-26.59** | **<0.001** |
| Stimulation | 0.00 | 0.09 | 66 | -0.05 | 0.96 |
| **Baseline RT** | **0.60** | **0.07** | **66** | **8.85** | **<0.001** |
| Block * Stimulation | 0.01 | 0.03 | 1186 | 0.47 | 0.64 |
| **Block \* Baseline Rt** | **-0.27** | **0.02** | **1186** | **-14.21** | **<0.001** |
| Stimulation * Baseline RT | 0.01 | 0.09 | 66 | 0.15 | 0.89 |
| **Block \* Stimulation \* Baseline RT** | **0.06** | **0.03** | **1186** | **2.16** | **0.03** |

**Table S3.** Results from the ordinal mixed-effects model regressing accuracy back on block number, stimulation group, and baseline accuracy.

| Predictor | b | S.E. | z | p |
| --- | --- | --- | --- | --- |
| **Block** | **0.42** | **0.10** | **4.43** | **<0.001** |
| Stimulation | 0.13 | 0.30 | 0.45 | 0.65 |
| **Baseline Accuracy** | **0.80** | **0.17** | **4.70** | **<0.001** |
| Block * Stimulation | 0.16 | 0.13 | 1.27 | 0.20 |
| Block * Baseline Accuracy | -0.04 | 0.08 | -0.50 | 0.62 |
| Stimulation * Baseline Accuracy | -0.02 | 0.33 | -0.07 | 0.95 |
| Block * Stimulation * Baseline Accuracy | -0.20 | 0.14 | -1.39 | 0.17 |

**Table S4.** Results from the mixed-effects model regressing participants self-reported retrieval back on block number, stimulation group, and baseline RT.

| Predictor | $\beta$ | S.E. | df | t | p |
| --- | --- | --- | --- | --- | --- |
| Intercept | -0.08 | 0.11 | 66 | -0.67 | 0.50 |
| **Block** | **0.63** | **0.02** | **1186** | **35.53** | **<0.001** |
| Stimulation | 0.14 | 0.16 | 66 | 0.90 | 0.37 |
| Baseline RT | -0.19 | 0.11 | 66 | -1.66 | 0.10 |
| Block * Stimulation | 0.02 | 0.02 | 1186 | 0.81 | 0.42 |
| Block * Baseline Rt | 0.00 | 0.02 | 1186 | -0.15 | 0.88 |
| Stimulation * Baseline RT | 0.04 | 0.16 | 66 | 0.24 | 0.82 |
| Block * Stimulation * Baseline RT | 0.00 | 0.02 | 1186 | -0.16 | 0.88 |

#### Theta/Beta Ratio results

**Table S5.** Results from the mixed-effects model regressing RT back on block number, stimulation, resting-state TBR, and baseline RT.

| Predictor | $\beta$ | S.E. | df | t | p |
| --- | --- | --- | --- | --- | --- |
| Intercept | 0.05 | 0.08 | 59 | 0.66 | 0.51 |
| **Block** | **-0.45** | **0.02** | **1084** | **-26.44** | **<0.001** |
| Stimulation | -0.04 | 0.10 | 59 | -0.42 | 0.68 |
| TBR (rest) | 0.04 | 0.08 | 59 | 0.58 | 0.57 |
| **Baseline RT** | **0.71** | **0.05** | **59** | **13.74** | **<0.001** |
| Block * Stimulation | -0.03 | 0.02 | 1084 | -1.46 | 0.14 |
| Block * TBR (rest) | -0.01 | 0.02 | 1084 | -0.85 | 0.40 |
| **Stimulation \* TBR (rest)** | **-0.21** | **0.10** | **59** | **-2.06** | **0.04** |
| Block * Stimulation * TBR (rest) | -0.02 | 0.02 | 1084 | -1.03 | 0.30 |

**Table S6.** Results from the mixed-effects model regressing RT back on block number, stimulation, online TBR, and baseline RT

| Predictor | $\beta$ | S.E. | df | t | p |
| --- | --- | --- | --- | --- | --- |
| Intercept | 0.03 | 0.07 | 56 | 0.40 | 0.69 |
| **Block** | **-0.45** | **0.02** | **1033** | **-26.50** | **<0.001** |
| Stimulation | 0.01 | 0.10 | 56 | 0.13 | 0.90 |
| TBR (online) | 0.01 | 0.06 | 56 | 0.08 | 0.94 |
| **Baseline RT** | **0.68** | **0.05** | **56** | **13.64** | **<0.001** |
| Block * Stimulation | -0.03 | 0.02 | 1033 | -1.40 | 0.16 |
| Block * TBR (Online) | -0.02 | 0.02 | 1033 | -1.26 | 0.21 |
| **Stimulation \* TBR (Online)** | **-0.25** | **0.10** | **56** | **-2.46** | **0.02** |
| Block * Stimulation * TBR (Online) | 0.02 | 0.02 | 1033 | 0.62 | 0.54 |

**Table S7.** Results from the ordinal mixed-effects model regressing accuracy back on block number, stimulation, resting-state TBR, and baseline accuracy.

| Predictor | b | S.E. | z | p |
| --- | --- | --- | --- | --- |
| **Block** | **0.41** | **0.10** | **4.07** | **<0.001** |
| Stimulation | -0.09 | 0.33 | -0.27 | 0.78 |
| TBR (rest) | 0.25 | 0.24 | 1.03 | 0.30 |
| **Baseline Accuracy** | **0.87** | **0.16** | **5.35** | **<0.001** |
| Block * Stimulation | 0.16 | 0.14 | 1.22 | 0.22 |
| Block * TBR (rest) | 0.01 | 0.10 | 0.10 | 0.92 |
| Stimulation * TBR (rest) | -0.29 | 0.32 | -0.90 | 0.37 |
| Block * Stimulation * TBR (rest) | -0.08 | 0.13 | -0.63 | 0.53 |

**Table S8.** Results from the ordinal mixed-effects model regressing accuracy back on block number, stimulation, online TBR, and baseline accuracy.

| Predictor | b | S.E. | z | p |
| --- | --- | --- | --- | --- |
| **Block** | **0.48** | **0.10** | **4.69** | **<0.001** |
| Stimulation | -0.15 | 0.35 | -0.42 | 0.68 |
| TBR (Online) | 0.17 | 0.22 | 0.78 | 0.44 |
| **Baseline Accuracy** | **0.86** | **0.16** | **5.22** | **<0.001** |
| Block * Stimulation | 0.06 | 0.14 | 0.45 | 0.66 |
| *Block \* TBR (Online)* | *0.17* | *0.09* | *1.80* | *0.07* |
| Stimulation * TBR (rest) | 0.12 | 0.34 | 0.37 | 0.71 |
| **Block \* Stimulation \* TBR (Online)** | **-0.33** | **0.15** | **-2.28** | **0.02** |

**Table S9.** Results from the mixed-effects model regressing participants self-reported retrieval back on block number, stimulation group, resting-state TBR, and baseline RT.

| Predictor | $\beta$ | S.E. | df | t | p |
| --- | --- | --- | --- | --- | --- |
| **Intercept** | **52.38** | **4.02** | **59** | **13.04** | **<0.001** |
| **Block** | **19.37** | **0.59** | **1084** | **32.81** | **<0.001** |
| Stimulation | 5.39 | 5.41 | 59 | 1.00 | 0.32 |
| TBR (rest) | -3.69 | 4.07 | 59 | -0.91 | 0.37 |
| **Baseline RT** | **-6.70** | **2.73** | **59** | **-2.45** | **0.02** |
| Block * Stimulation | 0.60 | 0.80 | 1084 | 0.76 | 0.45 |
| **Block \* TBR (rest)** | **-2.38** | **0.60** | **1084** | **-3.96** | **<0.001** |
| Stimulation * TBR (rest) | 5.62 | 5.42 | 59 | 1.04 | 0.30 |
| **Block \* Stimulation \* TBR (rest)** | **3.97** | **0.80** | **1084** | **4.97** | **<0.001** |

**Table S10.**
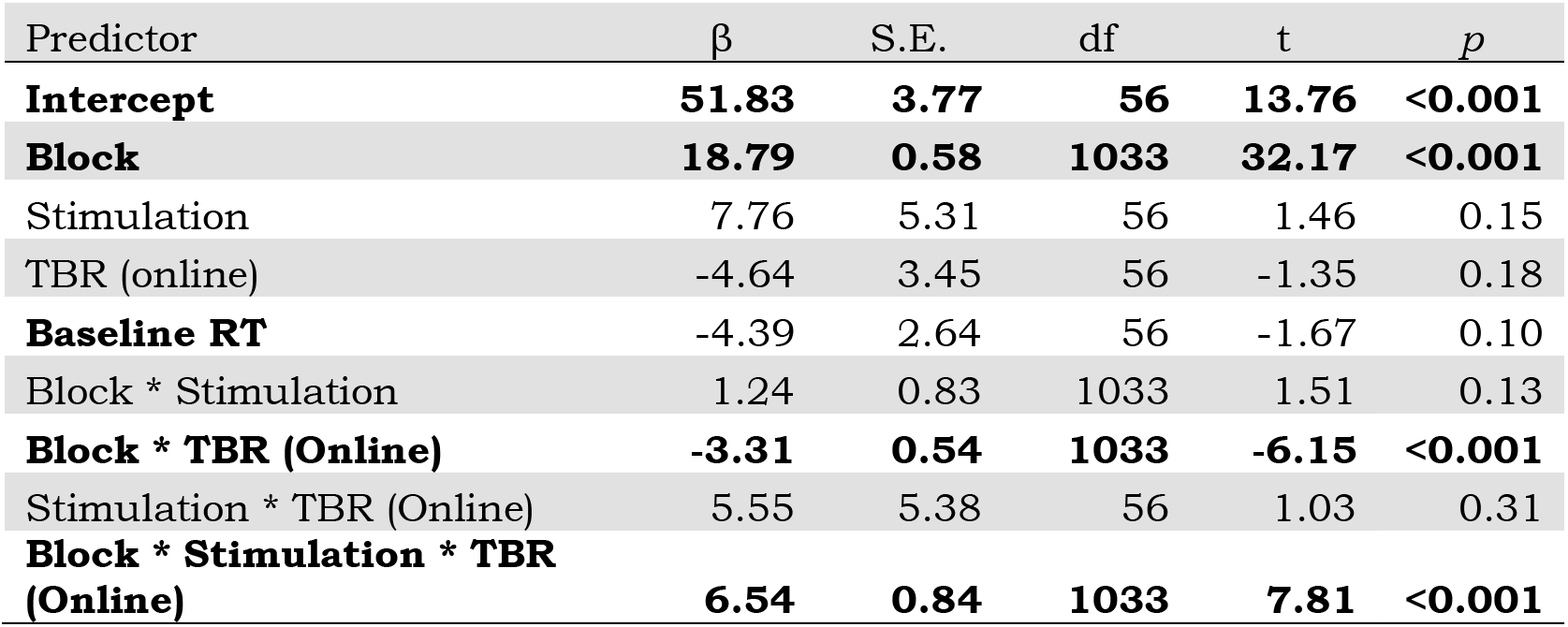
Results from the mixed-effects model regressing participants self-reported retrieval back on block number, stimulation group, online TBR, and baseline RT.

| Predictor | $\beta$ | S.E. | df | t | p |
| --- | --- | --- | --- | --- | --- |
| **Intercept** | **51.83** | **3.77** | **56** | **13.76** | **<0.001** |
| **Block** | **18.79** | **0.58** | **1033** | **32.17** | **<0.001** |
| Stimulation | 7.76 | 5.31 | 56 | 1.46 | 0.15 |
| TBR (online) | -4.64 | 3.45 | 56 | -1.35 | 0.18 |
| **Baseline RT** | **-4.39** | **2.64** | **56** | **-1.67** | **0.10** |
| Block * Stimulation | 1.24 | 0.83 | 1033 | 1.51 | 0.13 |
| **Block \* TBR (Online)** | **-3.31** | **0.54** | **1033** | **-6.15** | **<0.001** |
| Stimulation * TBR (Online) | 5.55 | 5.38 | 56 | 1.03 | 0.31 |
| **Block \* Stimulation \* TBR (Online)** | **6.54** | **0.84** | **1033** | **7.81** | **<0.001** |

#### Aperiodic results

**Table S11.** Results from the mixed-effects model regressing RT back on block number, stimulation, resting-state aperiodic exponent, and baseline RT

| Predictor | $\beta$ | S.E. | df | t | p |
| --- | --- | --- | --- | --- | --- |
| Intercept | 0.07 | 0.07 | 59 | 0.88 | 0.38 |
| **Block** | **-0.43** | **0.02** | **1084** | **-25.35** | **<0.001** |
| Stimulation | -0.05 | 0.10 | 59 | -0.46 | 0.65 |
| Aperiodic (rest) | 0.09 | 0.07 | 59 | 1.23 | 0.22 |
| **Baseline RT** | **0.72** | **0.05** | **59** | **14.19** | **<0.001** |
| **Block \* Stimulation** | **-0.04** | **0.02** | **1084** | **-1.97** | **0.05** |
| **Block \* Aperiodic (rest)** | **0.06** | **0.02** | **1084** | **3.32** | **<0.001** |
| **Stimulation \* Aperiodic (resting)** | **-0.29** | **0.10** | **59** | **-2.88** | **0.01** |
| **Block \* Stimulation \* Aperiodic (rest)** | **-0.12** | **0.02** | **1084** | **-5.09** | **<0.001** |

**Table S12.**
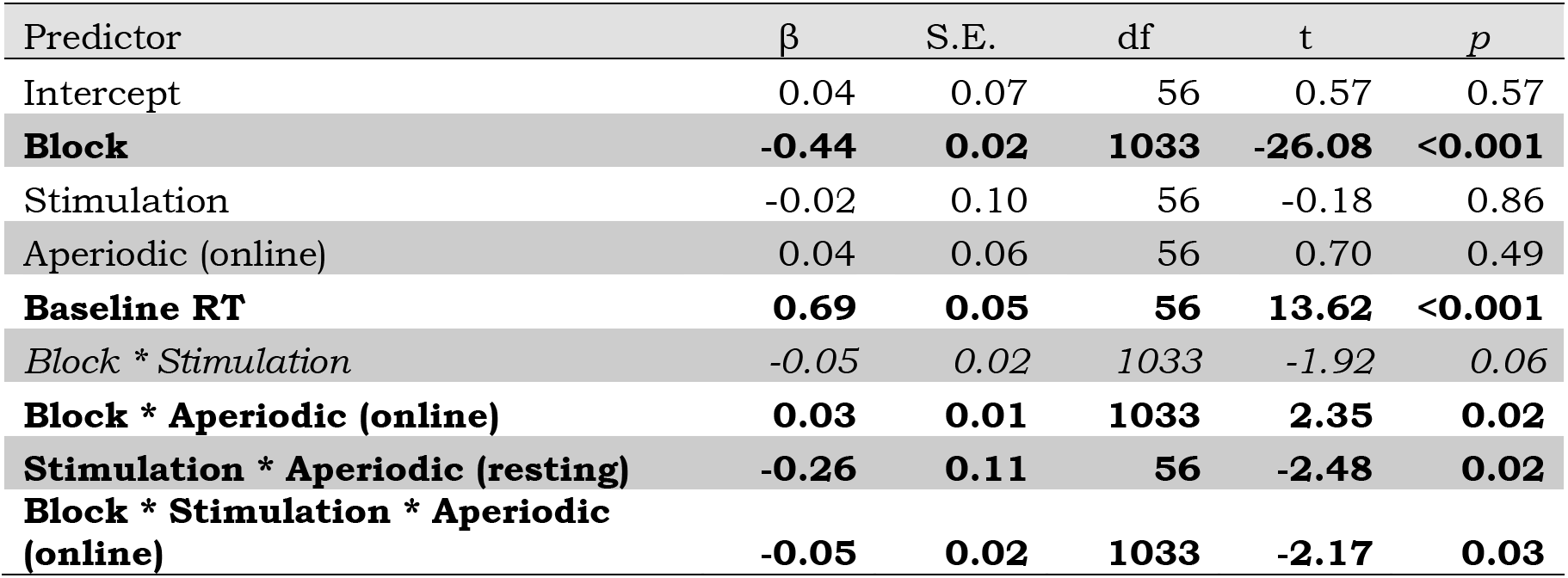
Results from the mixed-effects model regressing RT back on block number, stimulation, online aperiodic exponent, and baseline RT

**Table S13.**
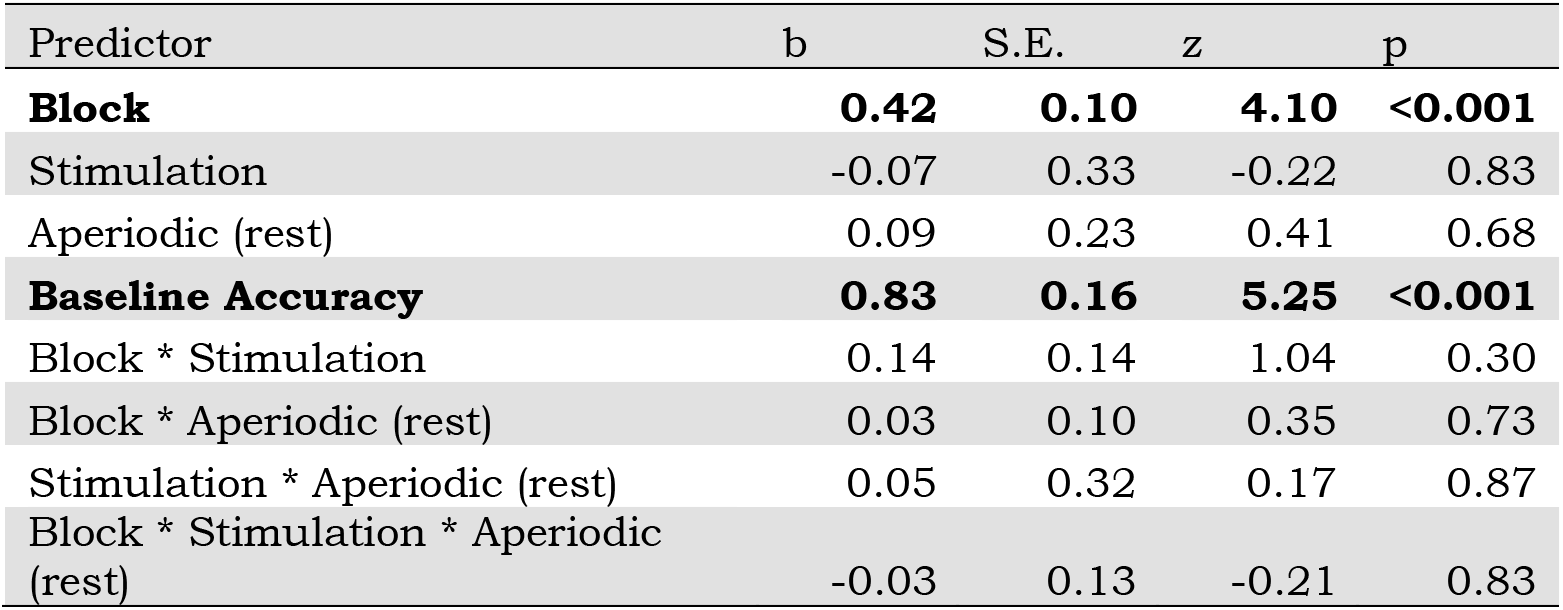
Results from the ordinal mixed-effects model regressing accuracy back on block number, stimulation, resting-state aperiodic exponent, and baseline accuracy.

**Table S14.** Results from the ordinal mixed-effects model regressing accuracy back on block number, stimulation, online aperiodic exponent, and baseline accuracy

| Predictor | b | S.E. | z | p |
| --- | --- | --- | --- | --- |
| **Block** | **0.46** | **0.10** | **4.68** | **<0.001** |
| Stimulation | -0.16 | 0.33 | -0.48 | 0.63 |
| Aperiodic (online) | 0.04 | 0.19 | 0.21 | 0.83 |
| **Baseline Accuracy** | **0.84** | **0.16** | **5.30** | **<0.001** |
| Block * Stimulation | 0.05 | 0.14 | 0.33 | 0.74 |
| **Block \* Aperiodic (online)** | **0.16** | **0.08** | **1.93** | **0.05** |
| **Stimulation \* Aperiodic (online)** | **0.69** | **0.33** | **2.12** | **0.03** |
| Block * Stimulation * Aperiodic (online) | -0.21 | 0.14 | -1.46 | 0.14 |

**Table S15.** Results from the mixed-effects model regressing participants self-reported retrieval back on block number, stimulation group, resting-state aperiodic exponent, and baseline RT.

| Predictor | $\beta$ | S.E. | df | t | p |
| --- | --- | --- | --- | --- | --- |
| **Intercept** | **52.15** | **4.00** | **59** | **13.05** | **<0.001** |
| **Block** | 19.03 | 0.59 | 1084 | 32.15 | **<0.001** |
| Stimulation | 4.88 | 5.37 | 59 | 0.91 | 0.37 |
| Aperiodic (rest) | -3.84 | 3.92 | 59 | -0.98 | 0.33 |
| **Baseline RT** | **-6.93** | **2.71** | **59** | **-2.56** | **0.01** |
| Block * Stimulation | 0.88 | 0.80 | 1084 | 1.11 | 0.27 |
| **Block \* Aperiodic (rest)** | **-3.27** | **0.58** | **1084** | **-5.62** | **<0.001** |
| Stimulation * Aperiodic (resting) | 8.84 | 5.35 | 59 | 1.65 | 0.10 |
| **Block \* Stimulation \* Aperiodic (rest)** | **4.92** | **0.79** | **1084** | **6.20** | **<0.001** |

**Table S16.** Results from the mixed-effects model regressing participants self-reported retrieval back on block number, stimulation group, online aperiodic exponent, and baseline RT.

| Predictor | $\beta$ | S.E. | df | t | p |
| --- | --- | --- | --- | --- | --- |
| **Intercept** | **51.95** | **3.69** | **56** | **14.09** | **<0.001** |
| **Block** | **18.89** | **0.57** | **1033** | **32.97** | **<0.001** |
| Stimulation | 8.20 | 5.20 | 56 | 1.58 | 0.12 |
| Aperiodic (online) | -5.12 | 3.21 | 56 | -1.59 | 0.12 |
| Baseline RT | -4.32 | 2.64 | 56 | -1.64 | 0.11 |
| Block * Stimulation | 1.33 | 0.81 | 1033 | 1.64 | 0.10 |
| **Block \* Aperiodic (online)** | **-3.54** | **0.50** | **1033** | **-7.06** | **<0.001** |
| Stimulation * Aperiodic (resting) | 3.94 | 5.44 | 56 | 0.72 | 0.47 |
| **Block \* Stimulation \* Aperiodic (online)** | **6.67** | **0.85** | **1033** | **7.88** | **<0.001** |

#### ERP findings

**Figure S2.**
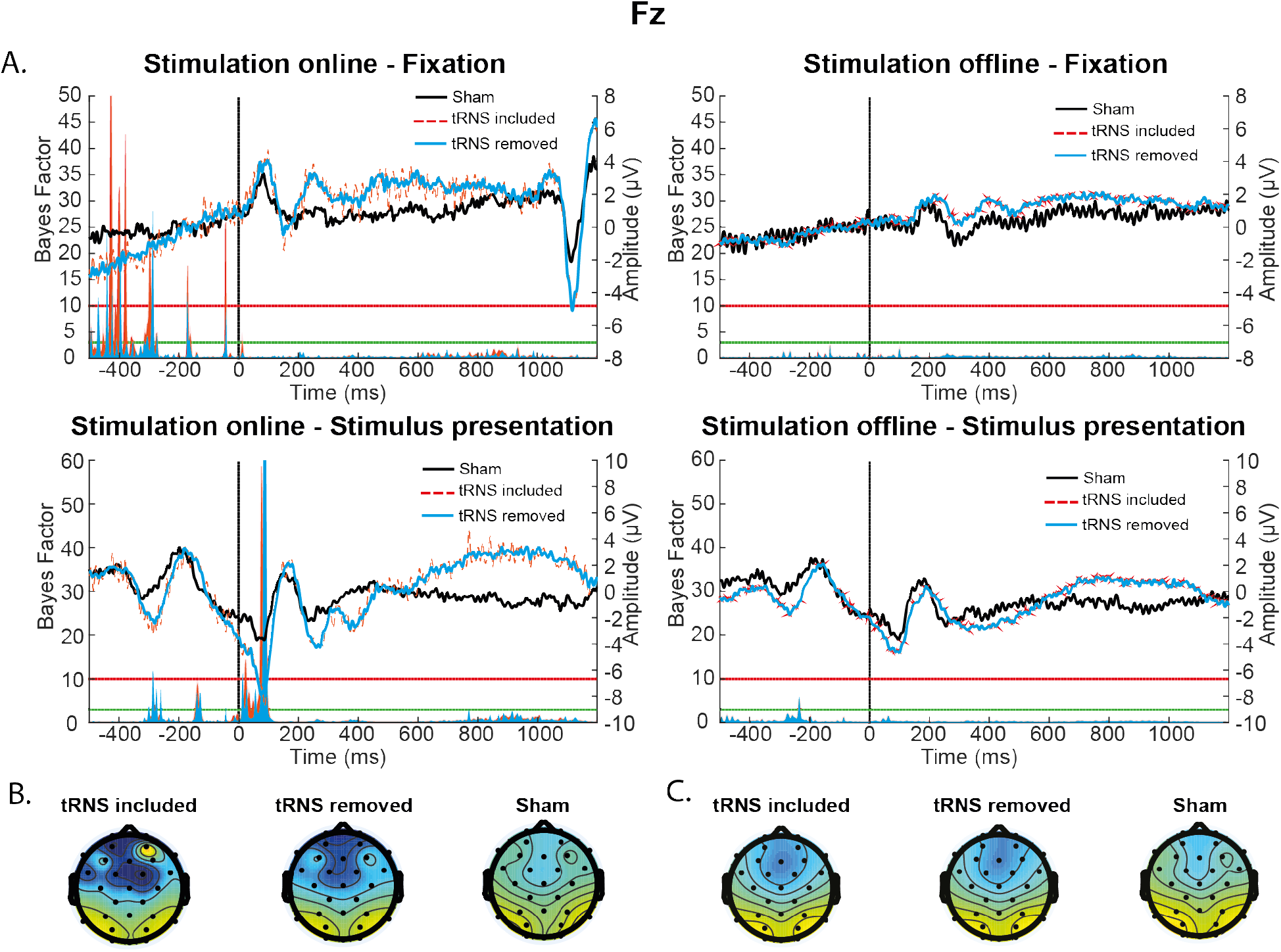
Task-related ERP effects at Fz following stimulus presentation. This figure is the same as Figure 10, but now also includes the data from the tRNS-included EEG A) ERPs following stimulus and fixation cross presentations showing a stimulation group effect during the online, but not the offline, period following stimulus presentation. Underneath the ERP trace plots, the BF comparisons of the tRNS removed and tRNS included groups against the sham group. They highlight differences in the ERP waveform between the tRNS and sham groups following stimulus presentation, primarily in early ERPs (~100ms following stimulus presentation). Topographic distribution of activity (in μV) for the sham and tRNS artefact removed data at the 100ms time point during the online (B) and offline period (C). The colour bar range for all topographies is the same.

**Figure S3.**
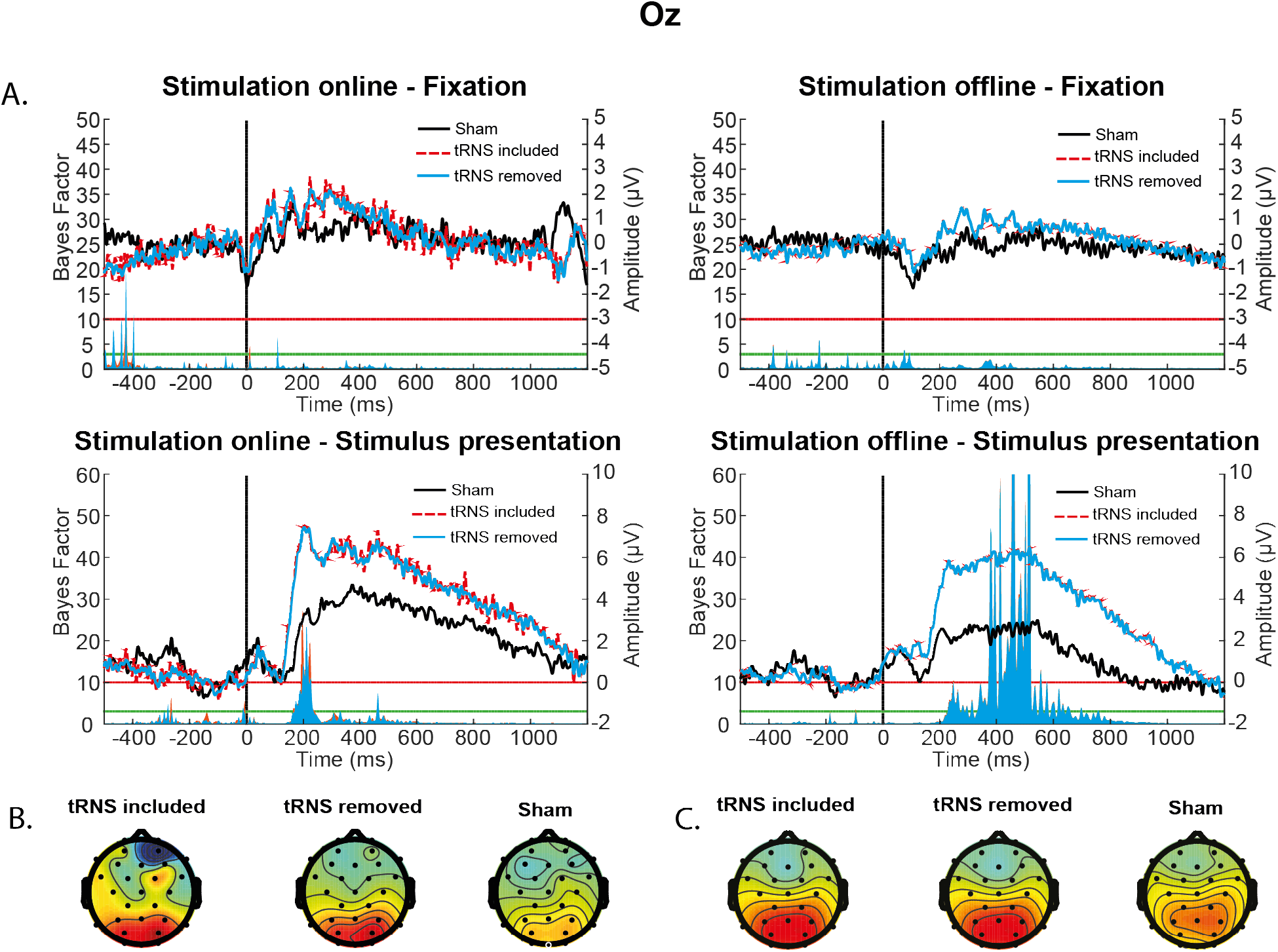
Task-related ERP effects at Oz following stimulus presentation. This figure is the same as Figure 11, but now also includes the data from the tRNS-included EEG. A) ERPs following stimulus and fixation cross presentations showing a stimulation group effect during the online, but not the offline, period following stimulus presentation. Underneath the ERP trace plots, the BF comparisons of the tRNS removed and tRNS included groups against the sham group. They highlight differences in the ERP waveform between the tRNS and sham groups following stimulus presentation, primarily in early ERPs (~100ms following stimulus presentation). Topographic distribution of activity (in μV) for the sham and tRNS artefact removed data at the 100ms time point during the online (B) and offline period (C). The colour bar range for all topographies is the same.

#### ERP multiple comparison

As a secondary test for group differences between sham and stimulation groups (following the removal of the stimulation artefact), we conducted a series of 1-DF tests similar to the methods outlined in Caspar et al. (2016). This method of analysis uses defined scaling factor for the prior distribution, which in this case was a normal distribution centred on 0, and a scaling factor defined by the grand average of the time window being analysed. Predefining a scaling factor based on the data itself offers two advantages over a more generic scaling factor such as that offered by Rouder et al. (2009). Firstly, the scaling factor can represent a difference or change in the data that could be reasonably expected of the task whilst Cohen’s d value of 0.7 suggested by Rouder et al. (2009) might be too large or small to be meaningful. Secondly, as discussed by Dienes (2019), for a meaningful application of Bayesian inference, our alternative models should have a defined theory behind them. If we know the tools with which we are measuring the indices of our theories, we should be able to approximately quantify a reasonable distribution for the effect. Specifically, as Dienes points out, as there is no default theory, there cannot be a default model.

In addition to calculating the BFs, we also calculated robustness regions (RR) for each analysis by finding the range in which the scaling factor for the prior lead to the same BF range. For example, if the BF for a comparison was above 10, the RR_BF_>10 would reflect the range of scaling factors which gave a BF greater than 10. The RR gives us an idea of how much the scaling factor is impacting our group differences

##### Fz results

Based on the grand average of all conditions (**Figure S4**), we noted 3 peaks of interest (−200–0ms; 50–120ms; 120-250ms), one of which was pre-stimulus. Our results indicated that the difference between sham and active stimulation groups was only noteworthy during the 50-120ms time window in the online period, with active tRNS was associated with a more negative N1 than the sham stimulation (−5.92μV vs. −2.74 μV, BF_N-3.9_=27.40, RR_BF>10_=[−14.65, −1.15]). Our results did not provide enough evidence for either hypotheses during the offline period (BF_N-3.9_=0.74, RR_0.3>BF>3_=[−10.15, 10.09]), and during the other two time windows (−200-0ms: BF_N-0.05_=0.91, RR_0.3>BF>3_=[−0.30,0.19], BF_N-0.05_=0.73, RR_0.3>BF>3_=[−0.30,0.19]; 120-250ms: BF_N-0.81_=0.95, RR_0.3>BF>3_=[−4.3, 4.1], BF_N-0.81_=0.92, RR_0.3>BF>3_=[−3.5, 3.4]),.

**Figure S4.**
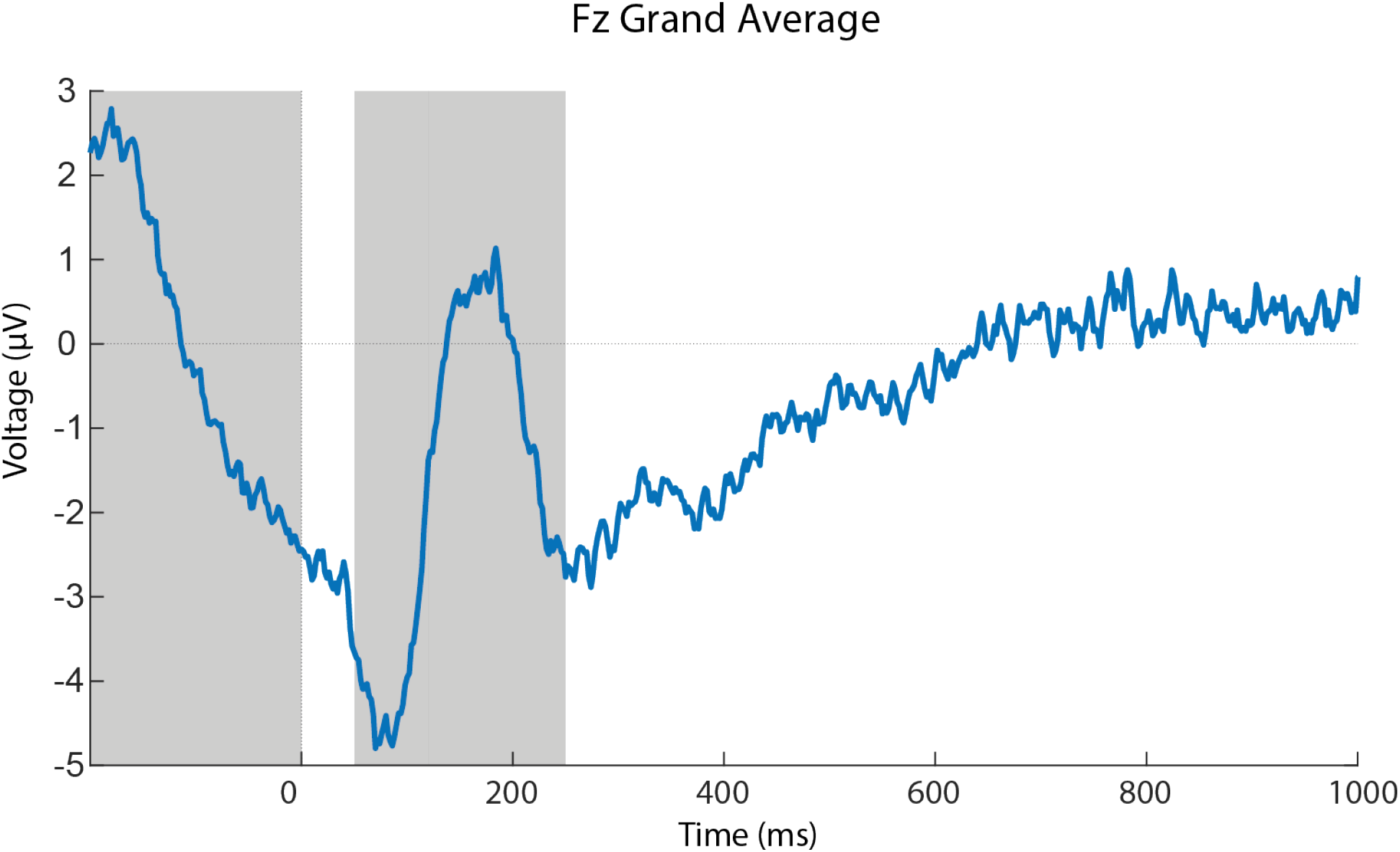
Grand average of ERP across all conditions taken from Fz. The grey windows reflect the peaks of interest.

##### Oz results

Based on the grand average of all conditions (**Figure S5**), we found 3 peaks of interest (0-100ms, 180-250ms, 280-600ms). For the 180-250ms peak, whilst stimulation was ongoing, there was a difference between the active and sham stimulation groups (BF_N 3.96_=29.55, RR_BF>10_=[1.46, 18.46]), with the active stimulation participants demonstrating a greater peak compared to sham stimulation (6.63μV vs. 2.80μV). This effect did not reach the threshold specified in the offline period (BF_N 3.96_=2.74, RR_0.3>BF>3_=[3.46, 28.96]). The late positive potential also demonstrated a significant difference between the two stimulation conditions, but this was in the offline period (BF_N 4.25_=54.61, RR_BF>10_=[−2.18, 29.25]), not the online period (BF_N4.25=_1.16, RR_0.3>BF>3_=[−18.74, 19.0]). Once again, the active stimulation group demonstrated a greater peak compared to the sham group (5.60μV vs. 2.27μV). By comparison, in the early peak (0-100ms, P1), there were no differences between the groups in either the online (BF_N 1.06_=0.50, RR_0.3>BF>3_=[−1.93, 1.81]) or offline (BF_N 1.06_=0.42, RR_0.3>BF>3_=[−1.43, 1.56]) period.

**Figure S5.**
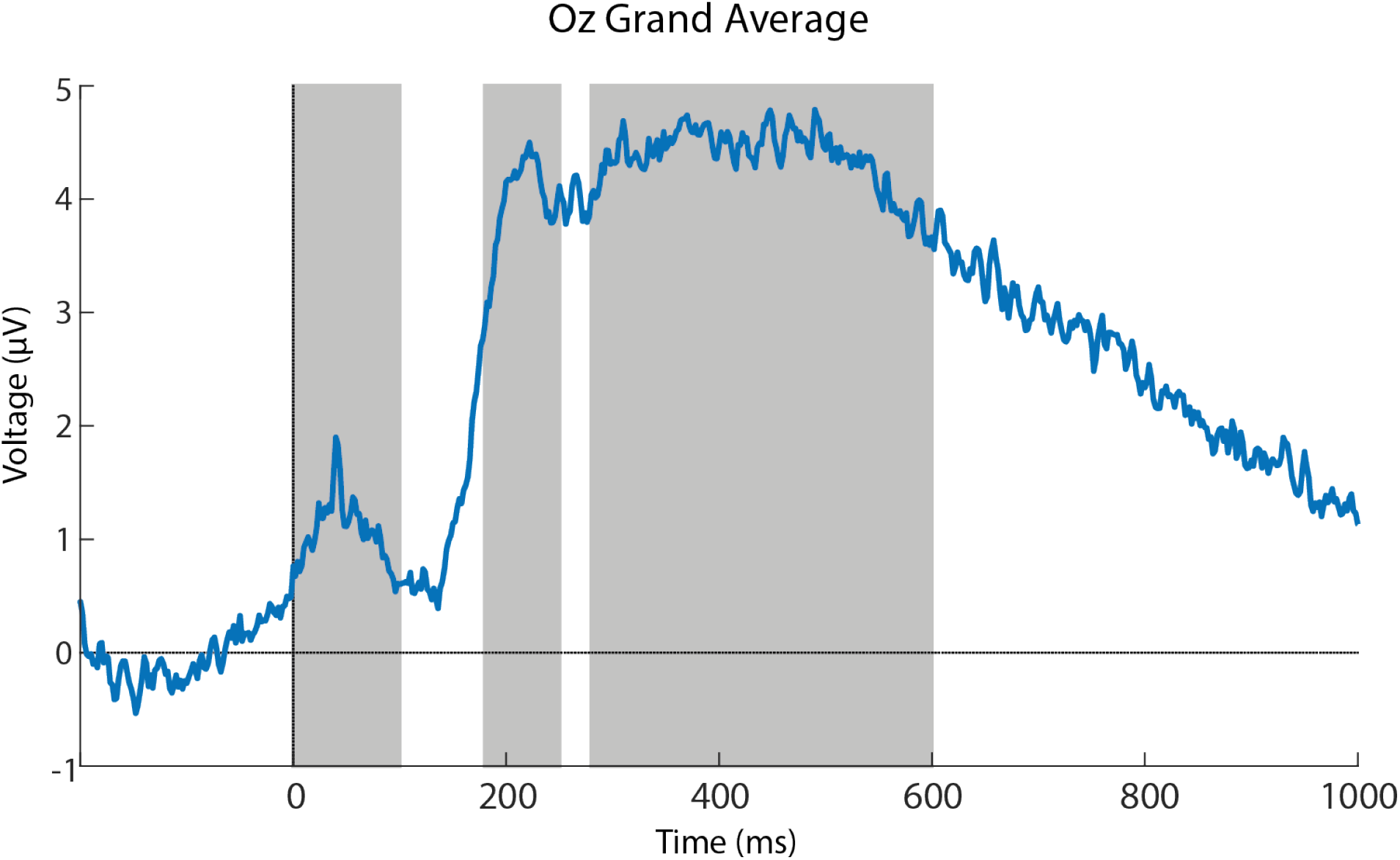
Grand average of ERP across all conditions taken from Oz. The grey windows reflect the peaks of interest.

##### Fixation results

All BF values for fixation results fell between 0.3 and 3, indicating that there is not enough evidence for either the alternative or null models.

#### ITPC results

Given previous research relating P2 amplitude to increased theta-band intertrial phase coherence (ITPC), we calculated the ITPC values for the theta band over Oz. These were then tested using Bayesian t-tests. The results indicated that whilst there were differences between active and sham tRNS groups in the P2 amplitude, there were no differences between active and sham tRNS groups in the theta-band ITPC over the same time period. There was, however, a difference between the tRNS artefact included conditions and both the sham group and tRNS removed condition during the online period. The tRNS artefact included data demonstrated lower ITPC values than both the sham and tRNS artefact removed data (BF_10_>10; **Figure S6**).

The theta ITPC values were reduced in the tRNS condition compared to the sham condition when the tRNS artefact remained in the data, suggesting that the increased P2 amplitude amongst the tRNS group was independent of increased phase locking as the P2 effect was seen in both the tRNS removed and the tRNS included datasets. In addition, the present results suggest that the association between P2 and ITPC, which has been based on correlational data, might not be due to the causal effect of P2 on ITPC. Future studies could shed further light on the association between P2 and ITPC and the potential causal effect of ITPC on P2.

**Figure S6:**
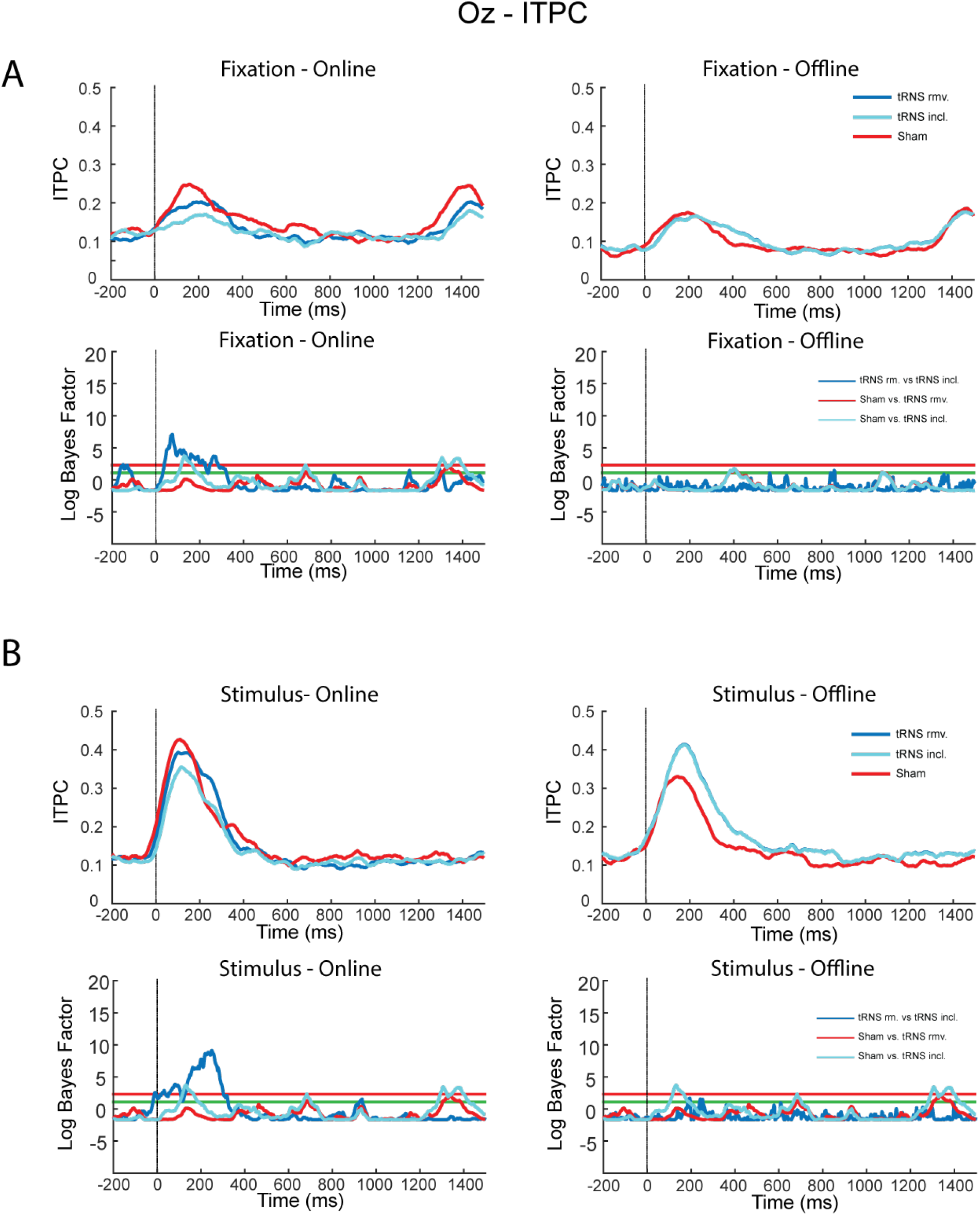
Changes in 6Hz ITPC at Oz following both fixation (panel A) and stimulus (panel B) presentations. Both fixation and stimulus presentations were associated with an increases in theta coherence across trials, peaking approximately 200ms after the stimulus event.

## Acknowledgements

We would like to thank both Prof. Zoltan Dienes and Prof. E.J. Wagenmakers for their insightful advice and discussion on Bayesian statistical analysis. We would also like to thank Makii Muthalib for his help. Finally, we would like to thank Dr. Jason Samaha and Dr. Franca Tecchio for their helpful comments on an earlier version of the manuscript. The Wellcome Centre for Integrative Neuroimaging is supported by core funding from the Wellcome Trust (203139/Z/16/Z). This work was supported by the European Research Council (Learning & Achievement 338065).

